# LSR Targets YAP to Modulate Intestinal Paneth Cell Differentiation

**DOI:** 10.1101/2022.10.30.514401

**Authors:** Yanan An, Chao Wang, Baozhen Fan, Ying Li, Feng Kong, Chengjun Zhou, Zhang Cao, Jieying Liu, Mingxia Wang, Hui Sun, Shengtian Zhao, Yongfeng Gong

## Abstract

Lipolysis-stimulated lipoprotein receptor (LSR) is a multi-functional protein that is best known for its roles in assembly of epithelial tricellular tight junctions and hepatic clearance of lipoproteins. Here, we investigated whether LSR contributes to intestinal epithelium homeostasis and pathogenesis of intestinal disease. By using multiple conditional deletion mouse models and *ex vivo* cultured organoids, we find that LSR elimination in intestinal stem cells results in disappearance of Paneth cell without affecting the differentiation of other cell lineages. Mechanistic studies reveal that LSR deficiency increases abundance and nuclear localization of YAP by modulating its phosphorylation and proteasomal degradation. Using gain- and loss-of-function studies we show that LSR protects against necrotizing enterocolitis through enhancement of Paneth cell differentiation in small intestinal epithelium. Thus, this study identifies LSR as an upstream negative regulator of YAP activity, an essential factor for Paneth cell differentiation, and a potential novel therapeutic target for inflammatory bowel disease.

## Introduction

Lipolysis-stimulated lipoprotein receptor (LSR), a multi-functional type I transmembrane protein containing an immunoglobulin-like domain (Masuda et al., 2011), has been linked to a variety of biological processes, molecular functions and cellular compartments. In the liver, LSR recognizes apolipoprotein B/E-containing lipoproteins in the presence of free fatty acids, and is thought to be involved in the hepatic clearance of triglyceride-rich lipoproteins (Narvekar et al., 2009; Yen et al., 1999). LSR also plays an essential role in organization of tricellular tight junctions that are involved in epithelial barrier function (Masuda et al., 2011; Sugawara et al., 2021). In the brain, LSR is specifically expressed at tricellular tight junctions between central nervous system endothelial cells and plays critical roles for proper blood–brain barrier formation and function (Sohet et al., 2015). LSR is also a target molecule for cell binding and internalization of Clostridium difficile transferase (Hemmasi et al., 2015; Papatheodorou et al., 2011). Although its role as either a tumor promoter or suppressor (or both) is not established, expression and localization of LSR have been found to be altered in several cancers (Dong et al., 2022; Shimada et al., 2017; Takahashi et al., 2021). The dominant subcellular localization of LSR is on the membrane, however, its localization in the nucleus of human epithelial cells has been reported (Reaves et al., 2017). Taken together, these findings suggest that the subcellular localization, function, and signaling pathways regulated by LSR are tissue- and cell type-specific. However, the specific cell types that express LSR have been difficult to identify, and the functions of LSR in postnatal development and tissue homeostasis have been hampered by the perinatal lethality of *Lsr* null mice (Mesli et al., 2004; Sohet et al., 2015).

Previous studies have revealed that LSR is highly expressed in the intestinal epithelium and localized at the basolateral membrane in addition to tricellular tight junctions of mouse intestine (Sugawara et al., 2021). However, roles of LSR in intestinal homeostasis remain to be fully elucidated. Earlier work in Drosophila has established a pivotal link between tricellular tight junction proteins, stem cell behaviour, and intestinal homeostasis (Resnik-Docampo et al., 2017), to what extent this connection is conserved in mammalian systems remains uncertain. Meanwhile, embryonic lethality in LSR-deficient mice highlights the importance of LSR for development (Mesli et al., 2004), but the significance and relevance of this protein in regulating intestinal development and differentiation in mammals are largely unclear. Here, we used a combination of *in vivo* conditional deletion mouse models and *ex vivo* cultured organoids to investigate the role of LSR in differentiation and function of intestinal epithelium.

## MATERIAL AND METHODS

The antibodies and primer sequences used for qRT-PCR used in this study are summarized in *SI Appendix*, Supplementary Table 1 and Supplementary Table 2, respectively. All mice were bred and maintained according to the Binzhou Medical University animal research requirements, and all procedures were approved by the Institutional Animal Research and Care committee. *SI Appendix, Materials and Methods* includes additional topics on generation of *Lsr* floxed animals, organoid culture, whole-transcriptome RNA sequencing, immunoprecipitation-mass spectrometry, co-immunoprecipitation, lentivirus-mediated knockdown, immunolabeling and confocal microscopy, statistical analyses, and so forth.

## Results

### LSR ablation results in increased numbers of proliferating cells and loss of Paneth cell lineage in the small intestine

To bypass the embryonic lethality of constitutive deletion and investigate the potential role of *Lsr* in intestinal homeostasis, we generated *Lsr^loxP/loxP^* mice, in which the mutant Lsr allele contains exons 1~2 flanked by loxP sites, on a C57BL/6J background (Figure S1A). We used *Lsr^loxP/loxP^; CAG-CreER* mice to achieve global *Lsr* deletion (Lsr^-/-^) upon tamoxifen treatment. We also generated intestinal-epithelium-specific Lsr-deficient mice (Lsr^vill KO^) by intercrossing villin-Cre and Lsr^loxP/loxP^ mice (Figure S1B). qRT-PCR and western blot analysis showed that the transcription and expression of LSR in the intestines of Lsr^vill KO^ and Lsr^-/-^ mice were successfully blocked (Figure 1A-D). LSR was expressed throughout the cellular membrane of epithelium in confocal sections, and did not show remarkable concentration at tricellular tight junctions in the Lsr^ctrl^ mouse intestine (Figure 1E). This was further ascertained by the disappearance of LSR signal in intestinal epithelium of Lsr^vill KO^ and Lsr^-/-^ mice (Figure 1E and F). In addition, transmission electron microscopy (TEM) showed that the structure of tight junctions between intestine epithelial cells (IECs) in Lsr^vill KO^ mice was not significantly different from that in Lsr^ctrl^ mice (Figure S1C). Further immunostaining of ZO-1 and CLDN1, as an indicator of bicellular tight junctions’ integrity, showed no difference between Lsr^vill KO^ and Lsr^ctrl^ small intestine (Figure S1D and E). The tricellular tight junctions’ integrity, assessed by tricellulin localization which was detected as dots at tricellular tight junctions, remained intact in Lsr^vill KO^ small intestine (Figure S1F). Together, these results indicate that the gross barrier function of tight junctions is not affected by deletion of LSR in IECs.

**Figure 1.**
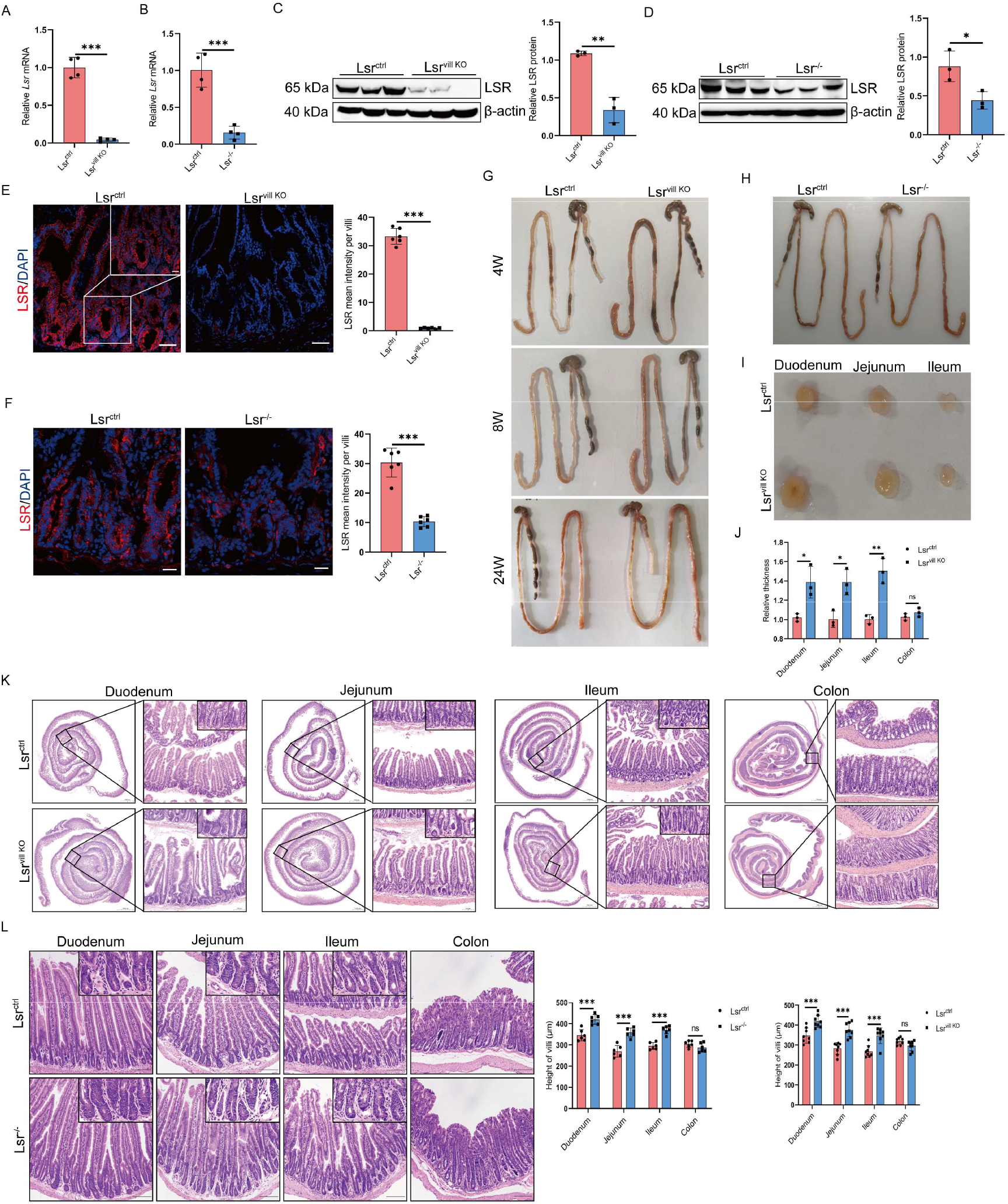
Generation and characterization of conditional Lsr knockout mice. (A and B) Expression of *Lsr* mRNA assessed by qRT-PCR in small intestines from Lsr^ctrl^ and Lsr^vill KO^ mice (A) and Lsr^ctrl^ and Lsr^-/-^ mice (B) (n=4). (C and D) Western blot and quantification analysis of LSR in small intestines from Lsr^ctrl^ and Lsr^vill KO^ mice (C) and Lsr^ctrl^ and Lsr^-/-^ mice (D) (n=3). (E and F) Immunofluorescence staining images and quantitative analysis of LSR in small intestines from Lsr^ctrl^ and Lsr^vill KO^ mice (E) and Lsr^ctrl^ and Lsr^-/-^ mice (F) (n=6). (G) Longitudinal views of intestine from Lsr^ctrl^ and Lsr^vill KO^ mice at 4, 8 and 24 weeks of age. (H) Longitudinal views of intestine from Lsr^ctrl^ and Lsr^-/-^ mice. (I) Transverse views of intestine from Lsr^ctrl^ and Lsr^vill KO^ littermate mice. (J) Analysis of intestinal wall thickness of Lsr^ctrl^ and Lsr^vill KO^ mice (n=3). (K) Representative H&E-stained transverse segments and villi length quantification of Lsr^ctrl^ and Lsr^vill KO^ mouse intestines (n=6). (L) Representative H&E-stained small intestine and villi length quantification of Lsr^ctrl^ and Lsr^-/-^ mice (n=6). Scale bars: E, 50 μm; F, 20 μm; K and J, 100 μm. *\*P < 0.05, **P < 0.01, ***P < 0.001*.

LSR deficiency had no effect on the overall structures of small intestine or colon (Figure 1G and H). Surprisingly, the small intestines of Lsr^vill KO^ mice were significantly thicker than those of Lsr^ctrl^ mice, and this phenomenon was consistent at various times during the growth of mice (Figure 1G, I and J). Moreover, we observed significantly longer villi in H&E-stained sections of small intestine of Lsr^vill KO^ and Lsr^-/-^ mice (Figure 1K and L). And expression of proliferation-related proteins Ki67 and proliferating cell nuclear antigen (PCNA) in the small intestine was found to be significantly increased in Lsr^vill KO^ mice (Figure 2A-D). Absence of LSR did not change the expression of OLFM4, a marker expressed by the Lgr5^+^ ISCs, in small intestine (Figure 2E and F; Figure S1G), indicating that the highly proliferating cells in Lsr^vill KO^ and Lsr^-/-^ mice were not ISCs. Since an increased percentage of Ki67 positive cells was predominantly found in the transit-amplifying zone (Figure 2A), we speculated that the proliferating cells may be undifferentiated progenitor cells. In summary, these data demonstrate that in the absence of LSR, ISCs do not contribute to the population of proliferating cells, presumably representing the transit-amplifying population leading to increased villus length.

**Figure 2.**
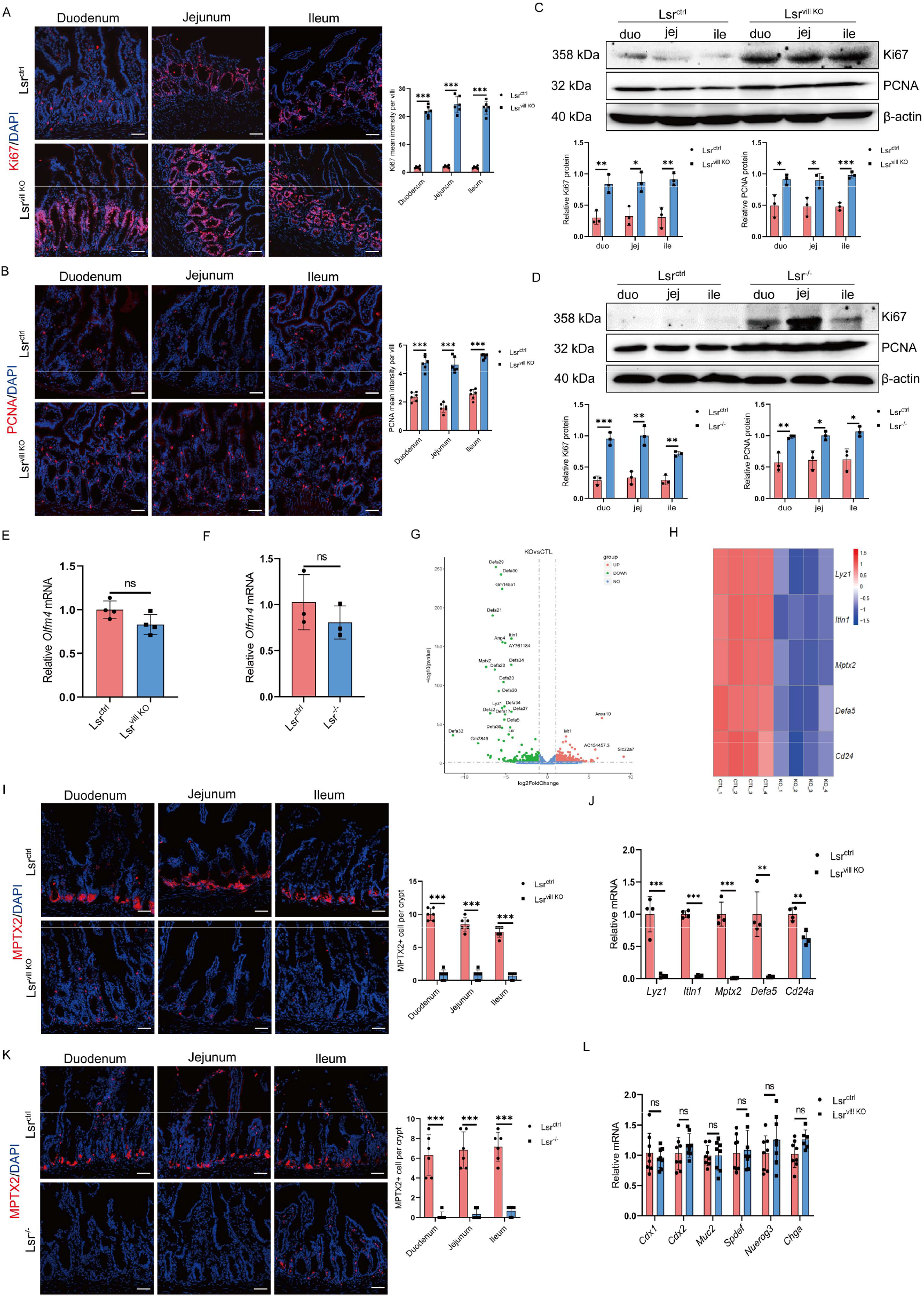
LSR ablation results in increased numbers of proliferating cells and loss of Paneth cell lineage in the small intestine. (A and B) Immunofluorescence staining images and quantitative analysis of Ki67 (A) and PCNA (B) in duodenal, jejunal and ileal segments from Lsr^ctrl^ and Lsr^vill KO^ mice (n=6). (C and D) Western blot and quantification analysis of Ki67 and PCNA protein expression in duodenal, jejunal and ileal segments from Lsr^ctrl^ and Lsr^vill KO^ mice (C) and Lsr^ctrl^ and Lsr^-/-^ mice (D) (n=3). (E and F) Expression of *Olfm4* mRNA assessed by qRT-PCR in small intestines from Lsr^ctrl^ and Lsr^vill KO^ mice (E) and Lsr^ctrl^ and Lsr^-/-^ mice (F) (n=4). (G) Differentially expressed genes (Lsr^vill KO^ vs. Lsr^ctrl^) are shown on a volcano plot (n=4; fold change > 2, *P < 0.05*). (H) Heatmap showing unsupervised hierarchical clustering of Paneth cell-related genes. (I) Immunofluorescence staining images and quantitative analysis of MPTX2 in duodenal, jejunal and ileal segments from Lsr^ctrl^ and Lsr^vill KO^ mice (n=6). (J) qRT-PCR validating several downregulated Paneth cell markers in small intestines from Lsr^ctrl^ and Lsr^vill KO^ mice (n=4). (K) Immunofluorescence staining images and quantitative analysis of MPTX2 in duodenal, jejunal and ileal segments from Lsr^ctrl^ and Lsr^-/-^ mice (n=6). (L) qRT-PCR analysis of the mRNA expression of columnar absorptive cell markers (*Cdx1* and *Cdx2*), goblet cell markers (*Muc2* and *Spdef*), and enteroendocrine cell markers (*Neurog3* and *Chga*) in small intestines from Lsr^ctrl^ and Lsr^vill KO^ mice (n=8). Scale bars: A, B, I and K, 50 μm. *\*P < 0.05, **P < 0.01, ***P < 0.001*.

In agreement with histological observations, which indicated the absence of granule-containing cells in H&E-stained sections of small intestine of Lsr^vill KO^ and Lsr^-/-^ mice (Figure 1K and L), RNA sequencing (RNA-seq) analysis of small intestinal lysate from Lsr^vill KO^ and Lsr^ctrl^ showed that Lsr-knockout downregulated many genes comprising the Paneth cell signature (*Lyz1, Itln1, Mptx2, Defa5*, and *Cd24*) (Figure 2G and H) and DEFA family members (Figure S1H). Data were validated by immunofluorescence (Figure 2I and K) and qRT-PCR (Figure 2J and Figure S1I) performed on the small intestine from Lsr^vill KO^, Lsr^-/-^ and control littermate mice. These results demonstrated that LSR elimination in intestinal epithelium leads to depletion of Paneth cells. Despite the absence of Paneth cells, expression levels of markers for other types of cells (EpCAM for IECs, MUC2 and SPDEF for goblet cells, CHGA and NEUROG3 for enteroendocrine cells, CDX1 and CDX2 for columnar absorptive cells, and LGR5 for stem cells) and Periodic acid–Schiff (PAS) staining of the goblet cells were not significantly changed (Figure 2L and Figure S1J-M). These data collectively suggest that LSR is absolutely required to maintain the Paneth cell lineage.

### LSR is required for Lgr5^+^ ISCs to Paneth cell differentiation

Paneth cells can be derived from Lgr5^+^ ISCs, we conjectured that LSR might affect the function of Lgr5^+^ ISCs to differentiate into Paneth cells. Interestingly, we found that LSR was highly expressed in Lgr5^+^ ISCs at the bottom of intestinal crypts (Figure 3A). We used a *Lgr5-CreERT2* strain to selectively delete *Lsr* in Lgr5^+^ ISCs in a tamoxifen-inducible manner (hereafter referred to as Lsr^ISC KO^) and to specifically explore the cellular and molecular consequences of *Lsr* deletion in Lgr5^+^ ISCs (Figure 3B and C). We found reduced expression of Paneth cell markers in the small intestine of Lsr^ISC KO^ mice 15 days after tamoxifen injection, again supporting the idea that LSR is required for Lgr5^+^ ISCs to Paneth cell differentiation (Figure 3D and E). In line with the findings in Lsr^vill KO^ mice, Lsr^ISC KO^ intestine possessed other intestinal cell types (Figure S2A and B). In Lsr^ISC KO^ mice, intestinal permeability assay showed that the concentration of FITC-Dextran 4000 in plasma seemed to increase compared to that of the Lsr^ctrl^ mice, but it did not attain a significant difference (Figure S2C), indicating the phenotype caused by LSR deficiency is not attributable to the dysfunction of permeability barrier. In contrast to Lsr^vill KO^ mice, we found no difference in the proliferative cells in small intestine of Lsr^ISC KO^ mice, indicating a potential direct role of LSR on transit-amplifying population. These results further demonstrated that in the absence of LSR in Lgr5^+^ ISCs, Paneth cells were not formed, but the differentiation of other IEC types was unaffected.

**Figure 3.**
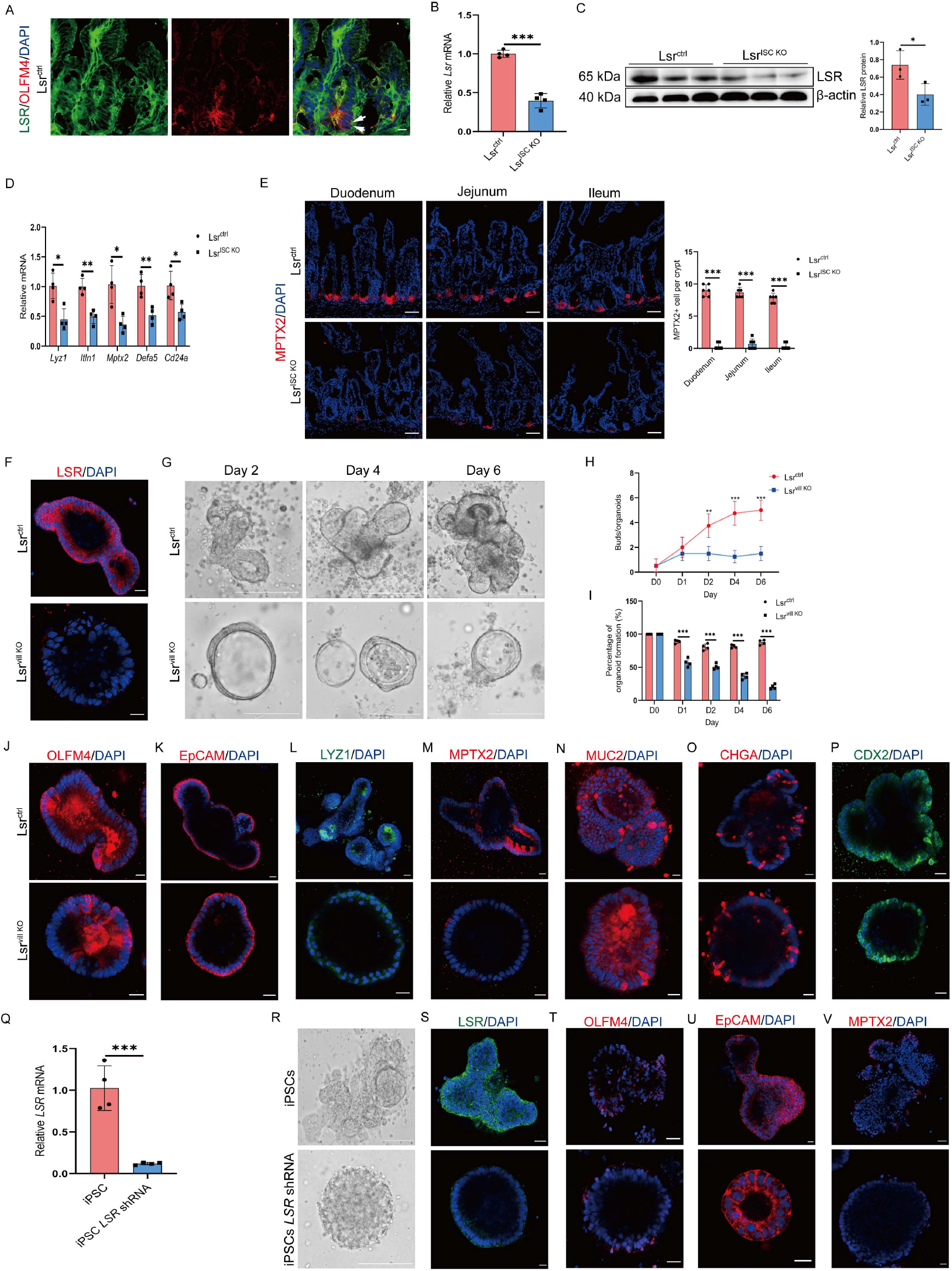
LSR is required for Lgr5^+^ ISCs to Paneth cell differentiation. (A) Immunofluorescence staining images of LSR and OLFM4 in small intestines from Lsr^ctrl^ mice (arrows indicated colocalization of LSR and OLFM4 in ISCs). (B) Expression of *Lsr* mRNA assessed by qRT-PCR in small intestines from Lsr^ctrl^ and Lsr^ISC KO^ mice (n=4). (C) Western blot and quantification analysis of LSR in small intestines from Lsr^ctrl^ and Lsr^ISC KO^ mice (n=3). (D) Expression of Paneth cell markers mRNA assessed by qRT-PCR in small intestines from Lsr^ctrl^ and Lsr^ISC KO^ mice (n=4). (E) Immunofluorescence staining images and quantitative analysis of MPTX2 in duodenal, jejunal and ileal segments from Lsr^ctrl^ and Lsr^ISC KO^ mice (n=6). (F and G) LSR immunofluorescence (F) and bright field images (G) of organoids established from intestinal crypts of Lsr^ctrl^ and Lsr^vill KO^ mice. (H and I) Quantification of the average number of buds per organoid (H) and percentage of organoid formation per well (n=3 wells per group) (I). (J-P) Immunofluorescence staining images OLFM4 (J), EpCAM (K), LYZ1 (L), MPTX2 (M), MUC2 (N), CHGA (O) and CDX2 (P) in organoids established from intestinal crypts of Lsr^ctrl^ and Lsr^vill KO^ mice. (Q) Expression of *LSR* mRNA assessed by qRT-PCR in control and *LSR* knockdown human iPSCs (n=4). (R-V) Bright field images (R) and immunofluorescence staining images of LSR (S) OLFM4 (T), EpCAM (U) and MPTX2 (V) in organoids differentiated from control and *LSR* knockdown human iPSCs. Scale bars: A, 10μm; E, S, T and V, 50 μm; F, J, K, L, M, N, O, P and U, 20 μm; G and R, 125 μm. *\*P < 0.05, **P < 0.01, ***P < 0.001*.

We also took advantage of crypt organoid culture to confirm the findings. Immunofluorescence analysis showed that LSR was expressed in organoid culture of Lsr^ctrl^ crypts and localized to the cell membrane, while no LSR expression was observed in Lsr^vill KO^ crypt organoids (Figure 3F). As expected, most crypts in Lsr^ctrl^ mice differentiated and budded into organoids with villi over time (Figure 3G). However, when intestinal crypts from Lsr^vill KO^ mice were cultured, *Lsr* deletion resulted in the formation of symmetrical spherical organoids (Figure 3F and G). In addition, on day 6, the shape of the spherical organoids did not change significantly; and the number of organoids gradually decreased (Figure 3G-I). Immunofluorescence staining showed that the distribution and quantity of stem cells (OFLM4) and epithelial cells (EpCAM) in Lsr^vill KO^ organoids were similar to those in Lsr^ctrl^ organoids (Figure 3J and K). Interestingly, similar to *in vivo* results, the Paneth cell markers (LYZ1 and MPTX2) exhibited significantly reduced and even complete loss of expression in Lsr^vill KO^ organoids (Figure 3L and M). Therefore, our results indicated that the Paneth cell number was positively correlated with the LSR expression level in both the crypt organoids and the intestine. Although the levels of the Paneth cell markers were significantly reduced in Lsr^vill KO^ organoids (Figure S2D), the numbers of other types of cells did not differ significantly between the Lsr^vill KO^ and Lsr^ctrl^ organoids (Figure 3N-P; Figure S2E). Therefore, the organoid culture of *Lsr* deficient crypts truthfully recapitulated the phenotype of Lsr^vill KO^ mice intestine. Next, we knocked down *LSR* in human iPSCs (Figure 3Q) and then cultured organoids. Unsurprisingly, we obtained results similar to those described above (Figure 3R-V), indicating a conserved function of LSR in the intestine from mouse to human. Collectively, these findings support a critical role for LSR in regulating ISCs differentiation to Paneth cells.

### Loss of LSR results in upregulation and activation of YAP

In intestine, excessive YAP can inhibit the differentiation of ISCs into Paneth cells (Gregorieff et al., 2015), and promote the proliferation of undifferentiated progenitor cells (Camargo et al., 2007). Recent work by Serra *et al*. revealed that homogeneous activation or suppression of YAP abolished Paneth cell differentiation and organoid budding, leading to the development of symmetrical spheres that were either develop into enterocysts or remain as undifferentiated (Serra et al., 2019). The loss of Paneth cells, proliferation of undifferentiated progenitor cells, and formation of symmetrical spherical organoids resulting from elimination of *Lsr* in the intestine led us to test whether abnormal YAP activation contributes to these phenotypes. Confocal imaging showed a colocalization of YAP with LSR in mouse intestine and Lgr5^+^ ISCs (Figure 4A), indicating that LSR might have a regulatory effect on YAP. Additionally, in Lsr^-/-^, Lsr^vill KO^ and Lsr^ISC KO^ mouse models, we observed enhanced YAP protein abundance accompanied by increased mRNA expression of the well-established YAP target genes, such as *Edn1, Ctgf*, and *Cyr61* (*Fan et al., 2022*), in the small intestine (Figure 4B-G), but the mRNA expression level of *Yap* did not change significantly (Figure 4E-G). To test directly whether elevated YAP contributes to the phenotype caused by LSR deletion, we lowered YAP signaling in Lsr^vill KO^ mice with verteporfin (VP), a pharmacologic inhibitor of YAP-TEAD association, twice per day from postnatal day 7 to day 21. The expression of MPTX2 was found to be significantly increased in small intestine of VP-treated Lsr^vill KO^ mice (Figure 4H). In addition, the mRNA levels of Paneth cell markers were significantly increased in VP-treated Lsr^vill KO^ mice (Figure S2F). These results indicated that VP treatment restored Paneth cell number. H&E staining showed that the small intestinal villi hyperproliferation were also partially suppressed in VP-treated Lsr^vill KO^ mice (Figure 4I). The expression of several YAP target genes was significantly suppressed by VP treatment (Figure 4J), indicating decreased YAP signaling in intestine of VP-treated Lsr^vill KO^ mice. Similar results were observed in Lsr^vill KO^ mice treated with lentiviruses carrying the *Yap* shRNA (Figure 4K-M; Figure S2G). These observations strongly support the causal link between the LSR-dependent YAP regulation and the altered Paneth cell differentiation in *Lsr*-null mice.

**Figure 4.**
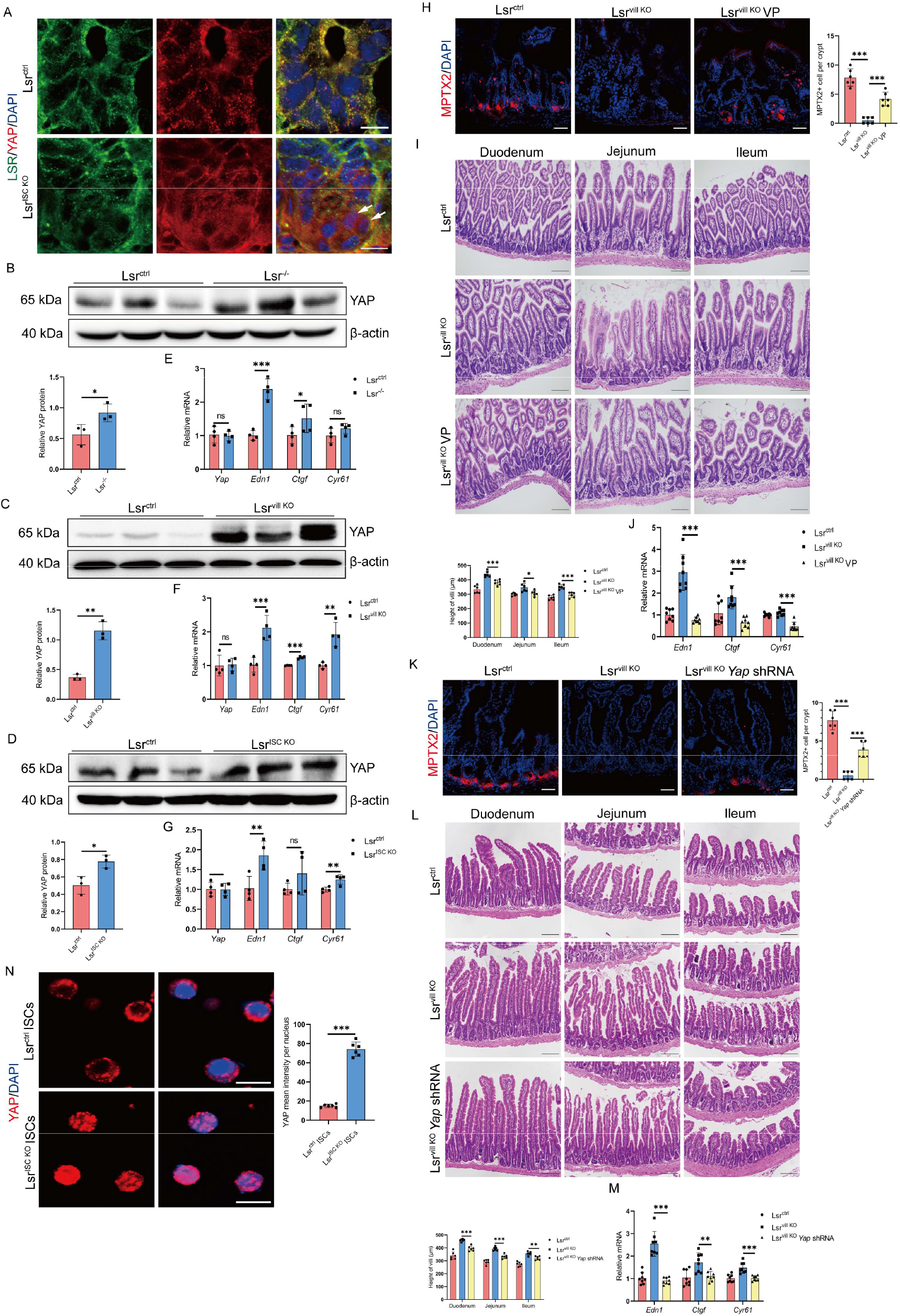
Loss of LSR results in upregulation and activation of YAP. (A) Immunofluorescence staining images of LSR and YAP in small intestines from Lsr^ctrl^ and Lsr^ISC KO^ mice (arrows indicated the ISCs). (B-D) Western blot and quantification analysis for YAP of small intestine lysates from Lsr^ctrl^ and Lsr^-/-^ mice (B), Lsr^ctrl^ and Lsr^vill KO^ mice (C) and Lsr^ctrl^ and Lsr^ISC KO^ mice (D) (n=3). (E-G) Expression of *Yap, Edn1, Ctgf*, and *Cyr61* mRNA assessed by qRT-PCR in small intestines from Lsr^ctrl^ and Lsr^-/-^ mice (E), Lsr^ctrl^ and Lsr^vill KO^ mice (F) and Lsr^ctrl^ and Lsr^ISC KO^ mice (G) (n=4). (H-J) Immunofluorescence staining images and quantitative analysis of MPTX2 (H), H&E staining and quantification analysis (I), and qRT-PCR analysis for YAP targets (J) in small intestines from Lsr^ctrl^ mice, Lsr^vill KO^ mice, and Lsr^vill KO^ mice treated with VP (n=6). (K-M) Immunofluorescence staining images and quantitative analysis of MPTX2 (K), H&E staining and quantification analysis (L) and qRT-PCR analysis for YAP targets (M) in small intestine from Lsr^ctrl^ mice, Lsr^vill KO^ mice, and Lsr^vill KO^ mice treated with lentivirus expressing *Yap* shRNA (n=8). (N) Immunofluorescence staining images and quantitative analysis of YAP in ISCs isolated from Lsr^ctrl^ and Lsr^ISC KO^ mice (n=6). Scale bars: A and N, 10 μm; H and K, 50 μm; I and L, 100 μm. *\*P < 0.05, **P < 0.01, ***P < 0.001*.

Next, we sought to verify the importance of YAP to the phenotype of *Lsr*-deficient intestine. We examined the expression of YAP in ISCs *in vitro*, and found that *Lsr* deletion led to significant increases in YAP expression and nuclear translocation in the Lsr^ISC KO^ (Figure 4N) and Lsr^vill KO^ ISCs (Figure S2H) compared to Lsr^ctrl^ ISCs. *Yap* shRNA lentiviral transduction reduced the levels of *Yap* down to 30% in organoids (Figure S2I), and rescued de novo crypt formation (Figure S2J) and normalized Paneth cell differentiation (Figure S2K). Taken together, these results confirmed a role for YAP activation in regulation of ISCs function downstream of LSR.

Wnt/β-catenin signaling pathway was discovered to be required for intestinal homeostasis and Paneth cell differentiation (Totaro et al., 2018). However, β-catenin of crypts in the Lsr^vill KO^ and Lsr^ISC KO^ mice was similar to that of control crypts (Figure S2L and M). Activation of YAP has been reported to directly inhibit Wnt signaling in the intestine (Cheung et al., 2020; Li et al., 2020), but LSR knockout did not affect expression of most of the β-catenin-target genes including *Ccnd1, Axin2*, and *Cd44* in both the intestinal tissue and Lgr5^+^ ISCs isolated from Lsr^vill KO^ and Lsr^ctrl^ mice (Figure S2N and O). This data may be due to alterations in negative feedback mediators in the Wnt pathway. Further, both ATOH1 and SOX9, two critical transcription factors for the differentiation of intestinal Paneth cell lineage, were not significantly altered by *Lsr* ablation in ISCs (Figure S2P-R).

### 14-3-3 zeta and PP2Ac are involved in the regulation of YAP by LSR

The above data clearly indicate a role of LSR on YAP expression and activation, we examined in closer detail whether Hippo pathway activity, which negatively regulates YAP, is perturbed by LSR ablation. Immunoblot analysis indicated that LSR ablation actually led to up-regulation of phosphorylated large tumor suppressor (p-Lats) (Figure S3A and B), a component of the mammalian Hippo pathway, which inhibits YAP nuclear translocation and promotes its proteasomal degradation. We conjectured that the elevation of p-Lats1 is induced by YAP activation via a potential negative feedback mechanism. Moreover, the increase of YAP protein level was not due to increased transcription, since *Yap* mRNA was slightly decreased in the small intestine (Figure 4F and G). We speculated that *Lsr* deletion may affect the metabolism of YAP and cause its protein level to increase, therefore, a protein synthesis inhibitor (cycloheximide, CHX) and a proteasome inhibitor (MG132) were used to determine the effect of LSR on YAP metabolism. The degradation rate of YAP in the Lsr^vill KO^ group was significantly lower than that in the Lsr^ctrl^ group after CHX treatment (Figure 5A). Moreover, MG132 blocked the turnover of YAP in the presence of CHX, and deletion of *Lsr* had an effect similar to that of MG132 on blocking YAP turnover (Figure 5B). Thus, deletion of LSR might increase YAP accumulation by suppressing its proteasomal degradation. Phosphorylation of YAP at Ser381 can increase its ubiquitination, ultimately leading to proteasomal degradation (Zhao et al., 2010). Decreased p-YAP (Ser381) (Figure 5C) and YAP ubiquitination (Figure 5D) were observed in the ISCs isolated from Lsr^vill KO^ mice. These results demonstrated that *Lsr* knockout enhanced YAP stability by decreasing its phosphorylation at Ser381, thereby blocking its ubiquitination and degradation.

**Figure 5.**
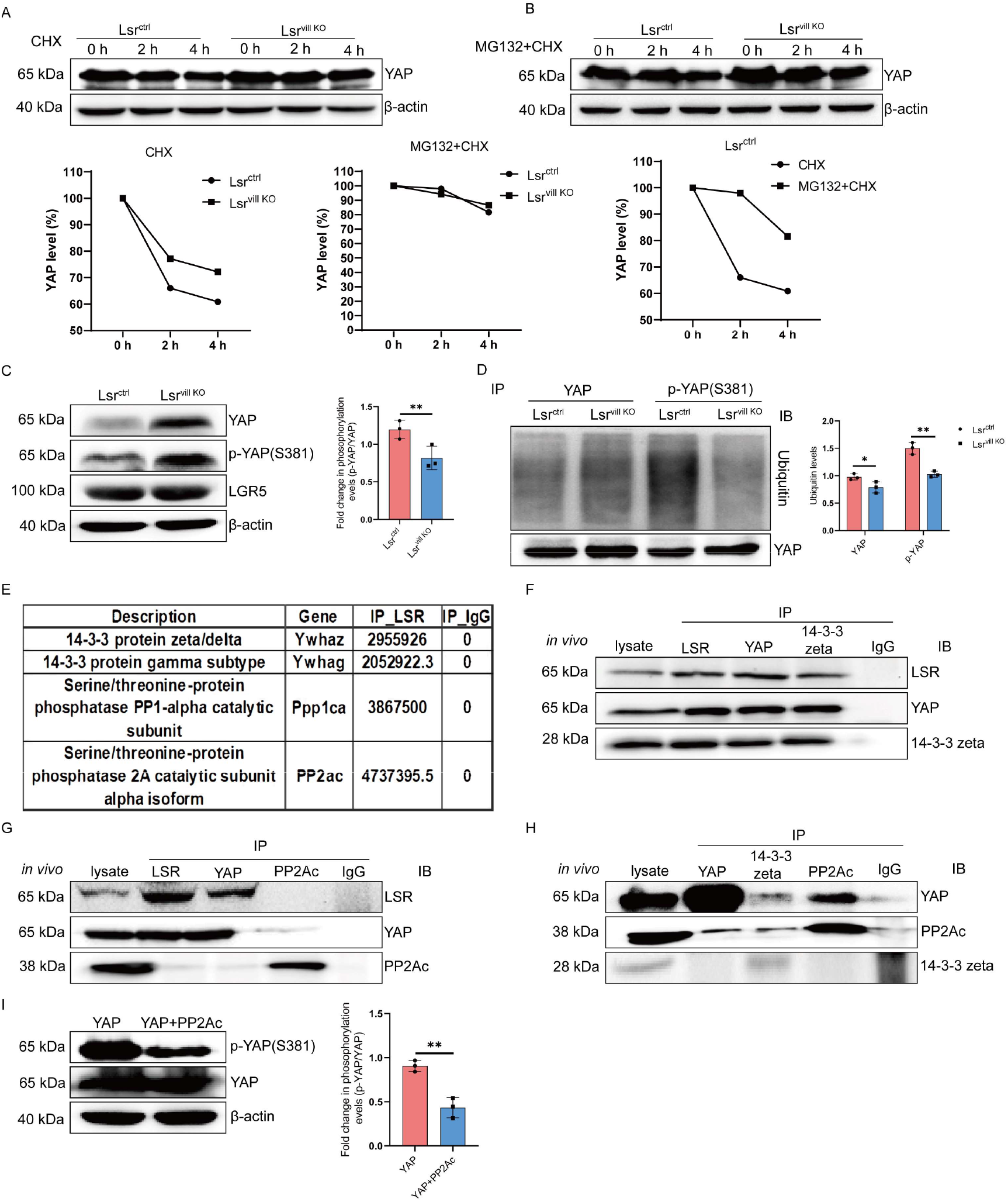
14-3-3 zeta and PP2Ac are involved in the regulation of YAP by LSR. (A and B) Western blot analysis for YAP in isolated ISCs treated with 5 mg/ml CHX for 0-4 h (A) or pretreated with 10 mM MG132 for 2 h and then treated with 5 mg/ml CHX for 0-4 h (B). And line graph showing the relative YAP protein levels quantified by YAP/β-actin ratio, which was arbitrarily set to 100% at the 0 hour point. (C) Western blot and quantification analysis for YAP, p-YAP (Ser381), LGR5 in isolated ISCs from Lsr^ctrl^ and Lsr^vill KO^ mice (n=3). (D) YAP ubiquitination levels and quantification analysis in isolated ISCs from Lsr^ctrl^ and Lsr^vill KO^ mice. (E) Mass spectrometry analysis of LSR immunocomplexes precipitated from isolated ISCs. (F) Co-IP showing that endogenous YAP, LSR, and 14-3-3 zeta interacts with each other in isolated ISCs from Lsr^ctrl^ mice. (G) Co-IP showing that endogenous YAP interacts with endogenous LSR, but not PP2Ac in isolated ISCs from Lsr^ctrl^ mice. (H) Co-IP showing that endogenous YAP interacts with endogenous PP2Ac, but not 14-3-3 zeta in isolated ISCs from Lsr^vill KO^ mice. (I) Western blot and quantification analysis of YAP and Ser381-phosphorylated YAP in HEK293 cells transfected YAP alone or together with PP2Ac (n=3). For all IP or Co-IP analysis, antibodies used for immunoprecipitation are shown above the lanes; antibody for blot visualization is shown on the right. *\*P < 0.05, **P < 0.01, ***P < 0.001*.

We next sought to determine the molecular mechanism through which LSR regulate the phosphorylation of YAP. We performed a mass spectrometry analysis of LSR immunocomplexes from ISCs. Results identified a list of 289 cellular proteins that specifically interact with LSR. We focused on two 14-3-3 proteins, 14-3-3 zeta and 14-3-3 gamma, and two protein phosphatases, protein phosphatase 1 catalytic subunit alpha (PPP1CA) and PP2Ac (Figure 5E), because they have been reported to participate in the regulation of YAP in multiple tissues (Hu et al., 2017; Meng et al., 2016; Schlegelmilch et al., 2011). It is also known that the turnover of YAP is regulated through its phosphorylation and ubiquitination in a 14-3-3 dependent manner. Remarkably, coimmunoprecipitation (co-IP) and immunoblot analysis of *in vivo* and *in vitro* samples confirmed that 14-3-3 zeta interacted with both YAP and LSR (Figure 5F, Figure S3C), but 14-3-3 gamma, PPP1CA, or PP2Ac did not interact with YAP or LSR either *in vivo* or *in vitro* (Figure S3D-H, Figure 5G). PPP1CA and PP2Ac have been reported to be phosphatases that directly and effectively dephosphorylate YAP (Ser381). An antagonistic and competitive interaction has been reported between PP2Ac and 14-3-3 for the phosphorylation of YAP (Schlegelmilch et al., 2011). Hence, we tested whether deletion of LSR results in an increased association of YAP with PP2Ac or PPP1CA. YAP interacted with PP2Ac instead of PPP1CA and exhibited reduced binding to 14-3-3 zeta in the ISCs from Lsr^vill KO^ mouse intestine (Figure 5H and Figure S3I) and HEK293 cells transfected with YAP, 14-3-3 zeta and PP2Ac expression plasmids (Figure S3J), while LSR and YAP did not seem to bind to PP2Ac in the ISCs from Lsr^ctrl^ mouse intestine or HEK293 cells (Figure 5G and Figure S3H). Therefore, YAP could interact with 14-3-3 zeta and PP2Ac respectively, and deletion of LSR might increase the interaction between YAP and PP2Ac by decreasing the YAP/14-3-3 zeta association. To determine whether PP2Ac can efficiently dephosphorylate YAP (Ser381), an *in vitro* phosphatase assay is performed in the presence of PP2Ac. The presence of PP2Ac resulted in diminished YAP (Ser381) phosphorylation and YAP accumulation (Figure 5I). These results suggested that loss of *Lsr* leads to more efficient association of YAP with PP2Ac, which can potentially dephosphorylate YAP at Ser381 and thereby decrease its degradation.

### Loss of *Lsr* exacerbates inflammation in a mouse model of NEC

Paneth cells, which are enriched in the ileum, have a central role in preventing intestinal inflammation (Adolph et al., 2013) and defects in Paneth cells are a hallmark of NEC. Infants who have developed NEC have decreased Paneth cell numbers compared to age-matched controls (Coutinho et al., 1998), and ablation of murine Paneth cells results in a NEC-like phenotype (Lueschow et al., 2018). We speculated that Paneth cell loss caused by LSR deletion may render the immature small intestine susceptible to development of NEC. Hence, to evaluate the role of LSR in NEC, we used 2,4,6-trinitrobenzenesulfonic acid (TNBS) to establish NEC models. Enteral administration of TNBS in 14-day-old mouse pups induced NEC, as revealed by villus disruption, clear separation of lamina propria and edema in the submucosa (Figure S4A and Figure 6A), increased mRNA levels of proinflammatory factors, including *Il-1α, Il-2, Il-6, Ifn-γ* and *Tnf-α* (Figure 6B), and increased infiltration of inflammatory cells (macrophages and neutrophil, Figure S4B and C, Figure 6C and D) in small intestine of mice treated with TNBS in Lsr^ctrl^ mice. As expected, LSR expression was significantly decreased in the small intestines of mice treated with TNBS (Figure 6E and F), and the number of Paneth cells was also significantly reduced (Figure S4D, Figure 6G and H). Compared with Lsr^ctrl^ mice, Lsr^vill KO^ mice showed worsening intestinal injury, represented by more severe transmural injury in the small intestine (Figure S4E and Figure 6I), increased mRNA levels of proinflammatory factors (Figure 6J), and increased infiltration of macrophages and neutrophils (Figure S4F and G, Figure 6K and L) after NEC induction. Collectively, these results implied that the lack of LSR led to more severe NEC. To further explore the effect of LSR on NEC, we established the NEC model in mice transduced with AAV encoding mouse *Lsr* (AAV-*Lsr*) or control vector (AAV-CTL). The LSR overexpression AAV decreased the severity of TNBS induced NEC (Figure S4H and Figure 6M), suppressed cytokines production (Figure 6N), decreased inflammatory cells infiltration (Figure S4 I and J; Figure 6O and P), and preserved Paneth cell number (Figure 6Q and R) in the small intestine of AAV-Lsr treated mice. These results suggest that LSR protects against TNBS induced NEC through upregulation of Paneth cell. To further verify the role of LSR in NEC, we tested intestinal samples from NEC patients. Compared with healthy control, patients with NEC exhibited severe intestinal injury (Figure 6S) and significantly reduced expression of LSR (Figure 6T), especially in ISCs (Figure 6U). Similar to previous reports (Underwood, 2012), infants with NEC have significantly decreased numbers of Paneth cells compared to age-matched controls (Figure 6V). These results confirmed that LSR plays an important role in the development of NEC and is expected to be a potential therapeutic target for NEC.

**Figure 6.**
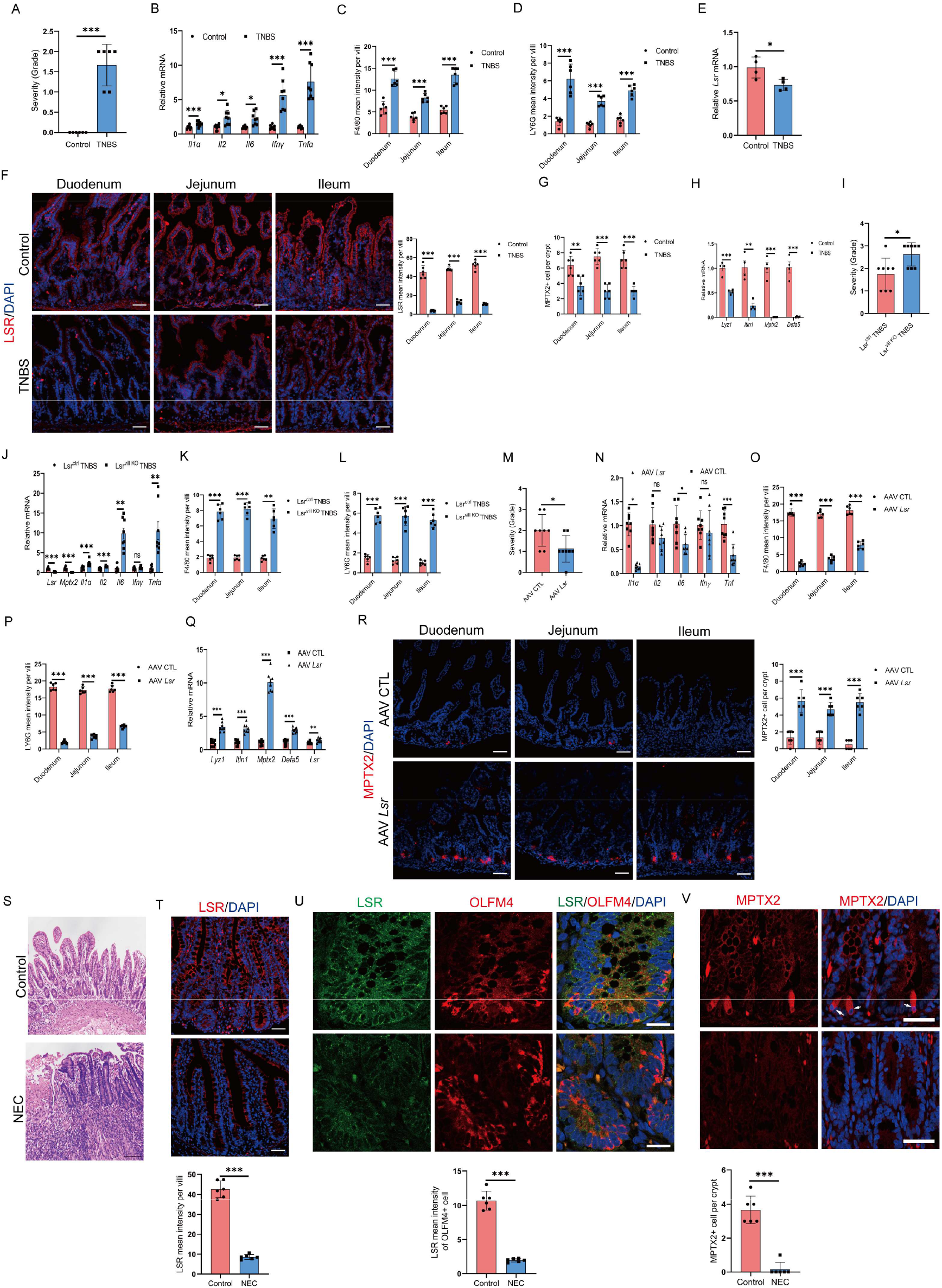
Loss of *Lsr* exacerbates inflammation in a mouse model of NEC. (A) Severity of intestinal injury in control and TNBS-treated mice (n=8). (B) Expression of *Il1α, Il2, Il6, Ifnγ* and *Tnfα* mRNA assessed by qRT-PCR in small intestines from control and TNBS-treated mice (n=8). (C and D) Quantification analysis of F4/80 (C) and LY6G (D) mean intensity per villi in small intestines from control and TNBS-treated mice (n=6). (E) Expression of *Lsr* mRNA assessed by qRT-PCR in small intestines from control and TNBS-treated mice (n=4). (F) Immunofluorescence staining images and quantitative analysis of LSR in small intestines from control and TNBS-treated mice (n=6). (G) Quantification analysis of MPTX2^+^ cell per crypt in small intestines from control and TNBS-treated mice (n=6). (H) mRNA expression of Paneth cell markers assessed by qRT-PCR in small intestines from control and TNBS-treated mice (n=4). (I) Severity of intestinal injury in TNBS-treated Lsr^ctrl^ and Lsr^vill KO^ mice (n=8). (J) mRNA expression of *Lsr, Mptx2, Il1α, Il2, Il6, Ifnγ* and *Tnfα* assessed by qRT-PCR in small intestines from TNBS-treated Lsr^ctrl^ and Lsr^vill KO^ mice (n=8). (K and L) Quantification analysis of F4/80 (K) and LY6G (L) mean intensity per villi in small intestines from TNBS-treated Lsr^ctrl^ and Lsr^vill KO^ mice (n=6). (M) Severity of intestinal injury in TNBS-treated mice administered with AAV-CTL or AAV-Lsr (n=8). (N) Expression of *Il1α, Il2, Il6, Ifnγ* and *Tnfa* mRNA assessed by qRT-PCR in small intestines from TNBS-treated mice administered with AAV-CTL or AAV-Lsr (n=8). (O and P) Quantification analysis of F4/80 (O) and LY6G (P) mean intensity per villi in small intestines from TNBS-treated mice administered with AAV-CTL or AAV-Lsr (n=6). (Q) Expression of *Lyz1, Itln1, Mptx2, Defa5* and *Lsr* mRNA assessed by qRT-PCR in small intestines from TNBS-treated mice administered with AAV-CTL or AAV-Lsr (n=8). (R) Immunofluorescence staining images and quantitative analysis of MPTX2 in small intestines from TNBS-treated mice administered with AAV-CTL or AAV-Lsr (n=6). (S) Representative H&E-stained small intestinal biopsy specimens of subjects with NEC and a control. (T-V) Immunofluorescence staining images and quantitative analysis of LSR (T), LSR and OLFM4 (U), and MPTX2 (V) in small intestinal biopsy specimens of subjects with NEC and a control (arrows indicated Paneth cells) (n=6). Scale bars: F, R, T and V, 50 μm; S, 100 μm; U, 20 μm. *\*P < 0.05, **P < 0.01, ***P < 0.001*.

## Discussion

Although the abundant expression of LSR has been described in intestine (Mesli et al., 2004; Sugawara et al., 2021), roles of this receptor in intestinal development, physiology and disease remain unclear. Here, we generated multiple conditional deletion mouse models to identify roles for LSR in the intestinal epithelium. We show that genetic ablation of *Lsr* in the intestinal epithelium results in increased numbers of proliferating cells, as well as a reduction in Paneth cell lineage. We demonstrate that the numbers of Lgr5^+^ ISCs quantified by OLFM4 immunostaining displayed no significant difference between Lsr^vill KO^ and control samples, suggesting that the associated increase in proliferating cells in Lsr^vill KO^ crypts is likely due to increased progenitor cell numbers. Strikingly, elimination of LSR from the intestinal epithelium or Lgr5^+^ ISCs resulted in almost complete disappearance of Paneth cell lineage, but had no effect on the number and distribution of other types of cells in small intestine. The disappearance of Paneth cell was reproduced in the tamoxifen-inducible Lsr^-/-^ and Lsr^ISC KO^ mice. An *ex vivo* organoid-forming assay showed that intestinal crypts from Lsr^vill KO^ mice or iPSCs with LSR knockdown formed symmetrical spherical organoids without Paneth cell lineage, implying that LSR plays a key role in the regulation of ISC function and differentiation toward Paneth cells in the intestinal epithelium.

YAP is a regulator of intestinal regeneration and ISCs fate (Gjorevski et al., 2022). We here show that the loss of LSR causes an increased abundance and enhanced nuclear translocation of YAP in Lsr^-/-^ and Lsr^vill KO^ intestinal epithelium and in Lsr^ISC KO^ ISCs. As for now, the molecular events and functional consequences that follow YAP activation remain controversial (Barry et al., 2013; Cai et al., 2010; Cheung et al., 2020; Gregorieff et al., 2015; Li et al., 2020; Zhou et al., 2011). Previous work has shown that overexpression of YAP results in enhanced proliferation of stem cell compartment and loss of differentiated cell types in small intestine (Camargo et al., 2007). Mice deficient for Hippo components Mst1 and Mst2 exhibited an expansion of stem-like undifferentiated cells and the almost complete disappearance of all secretory lineages, due to an increased abundance and enhanced nuclear translocation of YAP caused by the loss of Mst1/Mst2 (Zhou et al., 2011). Recent studies by several group have demonstrated that removal of Lats1/2, which caused the increase of YAP/TAZ expression and nuclear accumulation, resulted in expansion of the transit-amplifying population (Guillermin et al., 2021; Li et al., 2020). In agreement with these findings, the small intestine in Lsr^-/-^ and Lsr^vill KO^ mice was significantly thickened accompanied by increased number of transit-amplifying cells. However, genetic ablation of *Lsr* only resulted in loss of Paneth cell lineage, stem cells survived and produced various cell types of intestinal epithelium. Although Paneth cell development is tightly controlled by both Wnt and Notch pathways and their downstream transcription factors, recent studies pointed out that spatial heterogeneities in YAP activity are required for intestinal tissue morphogenesis, and homogeneous inhibitions or overexpression of YAP in all cells reduces Paneth cell differentiation (Gjorevski et al., 2022; Li, 2019). We confirm this finding and observe that homogeneous upregulation of YAP caused by LSR elimination from the intestinal epithelial compartment results in the almost complete disappearance of Paneth cell lineage.

Loss of Paneth cells was confirmed in organoid cultures derived from *Lsr* knockout mouse crypts or human iPSCs with stable *LSR* knockdown. Moreover, LSR deletion resulted in the formation of only symmetrical spherical organoids from mouse crypts or human iPSCs. These phenotypes are similar to YAP-overexpression organoids, where YAP is homogeneously active in all cells, and neither form Paneth cells nor display symmetry breaking (Serra et al., 2019). Originally, the transcriptional regulator YAP can stimulate single stem cells to enter a regenerative state and form a symmetric organoid (Serra et al., 2019); then, the emergence of Paneth cells can break the symmetry of the organoid, followed by of budding and the formation of villus structures (Chacon-Martinez et al., 2018; Sato et al., 2009). However, if YAP is overexpressed and localized in the nucleus, the symmetry cannot be broken, and the organoid remains spherical (Lukonin et al., 2020). Consistent with this concept, uniform induction in YAP activity, which also abrogated spatial gradients in YAP activity, resulted in *Lsr* knockout organoids remain symmetrical and contain no Paneth cells. Bearing in mind the pattern of YAP activity described above, we reasoned that uniform inhibition of YAP by treatment with *Yap* shRNA lentivirus could not restore the ability of *Lsr* knockout ISCs to differentiate into Paneth cells. Following YAP targeted knockdown, however, we observed rescued de novo crypt formation and enhanced Paneth cell number in *Lsr* mutant organoid. We do not currently know why such a difference exists. One possibility is that although lentiviral transduction is a highly efficient method to introduce targeted gene in organoids, the number of integrations per cell can be variable in organoids, resulting in heterogeneous knockdown efficiency of YAP within the cell population.

Next, we investigated the disease relevance of intestinal LSR using NEC as an intestinal injury model. Although NEC’s pathogenesis is multifactorial, Paneth cell depletion or dysfunction has been linked mechanistically to development of NEC. Our data clearly show that Paneth cell deficient mice caused by *Lsr* depletion displayed increased susceptibility to TNBS induced NEC. We found that the expression of LSR was significantly reduced in the intestines of NEC patients, and similar results were also obtained in NEC mouse models. Systemic delivery of LSR using an adenoviral delivery system profoundly inhibited the development and progression of NEC. These results reveal that LSR contributes to intestinal injury and disease progression in NEC, which will potentially advance our understanding of NEC, describe new biomarkers, and lead to novel therapeutic strategies for this multifactorial disease. It would be interesting to investigate the contribution of LSR to other gut inflammatory disorders such as Crohn’s disease apart from NEC, because misfunctioning Paneth cells accelerate the progress of these disorders (Wehkamp and Stange, 2020).

YAP signaling activity can be regulated by multiple factors, however, extracellular ligands, cell surface receptors, and signaling pathways regulating YAP have not been thoroughly examined. Our results demonstrate that LSR can directly impact the differentiation of ISCs into Paneth cells via regulating the degradation of YAP in the ISCs (Figure S5). This study establishes an important regulatory role of LSR in restricting YAP activity, however, many questions remain. Are there additional direct or indirect effects of LSR on YAP signaling activity? Are these tissue-specific or universal? Whether LSR can be used to combat excess YAP activity? Can LSR also act independently of the YAP pathway to control intestinal epithelium homeostasis? Intense studies are currently underway in our laboratory to address those important questions.

## Author contributions

Y. A., C. W., and B. F. designed and conducted *in vivo* and *in vitro* experiments, performed data analysis, and wrote the manuscript. Y. L., F. K., C. Z. and Z. C. performed histologic analysis. J. L., M. W. and H. S. performed mice genotyping. Y. G. and S. Z. designed and jointly supervised the study, analyzed the data, and wrote the manuscript.

## Acknowledgments

We thank Ping Li for animal husbandry. We thank the electron microscopic core lab of Shandong University for assistance on electron microscopy imaging. The work is funded by National Natural Science Foundation of China (81670620, 81870485, 81970578, 81971281 and 32200758), Taishan Scholars Program of Shandong Province (ts20190953), Natural Science Foundation of Shandong Province (ZR2020QH066, ZR2021QH115 and ZR2021QC105), Shandong First Medical University Academic Promotion Program (2020LI001), and Binzhou Medical University Independent Research Program.

## References

Adolph, T.E., Tomczak, M.F., Niederreiter, L., Ko, H.J., Bock, J., Martinez-Naves, E., Glickman, J.N., Tschurtschenthaler, M., Hartwig, J., Hosomi, S., et al. (2013). Paneth cells as a site of origin for intestinal inflammation. Nature 503, 272–276.

Barry, E.R., Morikawa, T., Butler, B.L., Shrestha, K., de la Rosa, R., Yan, K.S., Fuchs, C.S., Magness, S.T., Smits, R., Ogino, S., et al. (2013). Restriction of intestinal stem cell expansion and the regenerative response by YAP. Nature 493, 106–110.

Cai, J., Zhang, N., Zheng, Y., de Wilde, R.F., Maitra, A., and Pan, D. (2010). The Hippo signaling pathway restricts the oncogenic potential of an intestinal regeneration program. Genes & development 24, 2383–2388.

Camargo, F.D., Gokhale, S., Johnnidis, J.B., Fu, D., Bell, G.W., Jaenisch, R., and Brummelkamp, T.R. (2007). YAP1 increases organ size and expands undifferentiated progenitor cells. Current biology: CB 17, 2054–2060.

Chacon-Martinez, C.A., Koester, J., and Wickstrom, S.A. (2018). Signaling in the stem cell niche: regulating cell fate, function and plasticity. Development 145.

Cheung, P., Xiol, J., Dill, M.T., Yuan, W.C., Panero, R., Roper, J., Osorio, F.G., Maglic, D., Li, Q., Gurung, B., et al. (2020). Regenerative Reprogramming of the Intestinal Stem Cell State via Hippo Signaling Suppresses Metastatic Colorectal Cancer. Cell stem cell 27, 590–604 e599.

Coutinho, H.B., da Mota, H.C., Coutinho, V.B., Robalinho, T.I., Furtado, A.F., Walker, E., King, G., Mahida, Y.R., Sewell, H.F., and Wakelin, D. (1998). Absence of lysozyme (muramidase) in the intestinal Paneth cells of newborn infants with necrotising enterocolitis. Journal of clinical pathology 51, 512–514.

Dong, X., Zhang, X.B., Liu, P., Tian, Y., Li, L., and Gong, P. (2022). Lipolysis-Stimulated Lipoprotein Receptor Impairs Hepatocellular Carcinoma and Inhibits the Oncogenic Activity of YAP1 via PPPY Motif. Front Oncol 12.

Fan, S., Gao, Y., Qu, A., Jiang, Y., Li, H., Xie, G., Yao, X., Yang, X., Zhu, S., Yagai, T., et al. (2022). YAP-TEAD mediates PPAR alpha-induced hepatomegaly and liver regeneration in mice. Hepatology 75, 74–88.

Gjorevski, N., Nikolaev, M., Brown, T.E., Mitrofanova, O., Brandenberg, N., DelRio, F.W., Yavitt, F.M., Liberali, P., Anseth, K.S., and Lutolf, M.P. (2022). Tissue geometry drives deterministic organoid patterning. Science 375, eaaw9021.

Gregorieff, A., Liu, Y., Inanlou, M.R., Khomchuk, Y., and Wrana, J.L. (2015). Yap-dependent reprogramming of Lgr5(+) stem cells drives intestinal regeneration and cancer. Nature 526, 715–718.

Guillermin, O., Angelis, N., Sidor, C.M., Ridgway, R., Baulies, A., Kucharska, A., Antas, P., Rose, M.R., Cordero, J., Sansom, O., et al. (2021). Wnt and Src signals converge on YAP-TEAD to drive intestinal regeneration. The EMBO journal 40, e105770.

Hemmasi, S., Czulkies, B.A., Schorch, B., Veit, A., Aktories, K., and Papatheodorou, P. (2015). Interaction of the Clostridium difficile Binary Toxin CDT and Its Host Cell Receptor, Lipolysis-stimulated Lipoprotein Receptor (LSR). The Journal of biological chemistry 290, 14031–14044.

Hu, J.K., Du, W., Shelton, S.J., Oldham, M.C., DiPersio, C.M., and Klein, O.D. (2017). An FAK-YAP-mTOR Signaling Axis Regulates Stem Cell-Based Tissue Renewal in Mice. Cell stem cell 21, 91–106 e106.

Li, Q., Sun, Y., Jarugumilli, G.K., Liu, S., Dang, K., Cotton, J.L., Xiol, J., Chan, P.Y., DeRan, M., Ma, L., et al. (2020). Lats1/2 Sustain Intestinal Stem Cells and Wnt Activation through TEAD-Dependent and Independent Transcription. Cell stem cell 26, 675–692 e678.

Li, V.S.W. (2019). Yap in regeneration and symmetry breaking. Nature cell biology 21, 665–667.

Lueschow, S.R., Stumphy, J., Gong, H., Kern, S.L., Elgin, T.G., Underwood, M.A., Kalanetra, K.M., Mills, D.A., Wong, M.H., Meyerholz, D.K., et al. (2018). Loss of murine Paneth cell function alters the immature intestinal microbiome and mimics changes seen in neonatal necrotizing enterocolitis. PloS one 13, e0204967.

Lukonin, I., Serra, D., Challet Meylan, L., Volkmann, K., Baaten, J., Zhao, R., Meeusen, S., Colman, K., Maurer, F., Stadler, M.B., et al. (2020). Phenotypic landscape of intestinal organoid regeneration. Nature 586, 275–280.

Masuda, S., Oda, Y., Sasaki, H., Ikenouchi, J., Higashi, T., Akashi, M., Nishi, E., and Furuse, M. (2011). LSR defines cell corners for tricellular tight junction formation in epithelial cells. Journal of cell science 124, 548–555.

Meng, Z., Moroishi, T., and Guan, K.L. (2016). Mechanisms of Hippo pathway regulation. Genes & development 30, 1–17.

Mesli, S., Javorschi, S., Berard, A.M., Landry, M., Priddle, H., Kivlichan, D., Smith, A.J., Yen, F.T., Bihain, B.E., and Darmon, M. (2004). Distribution of the lipolysis stimulated receptor in adult and embryonic murine tissues and lethality of LSR-/-embryos at 12.5 to 14.5 days of gestation. European journal of biochemistry 271, 3103–3114.

Narvekar, P., Berriel Diaz, M., Krones-Herzig, A., Hardeland, U., Strzoda, D., Stohr, S., Frohme, M., and Herzig, S. (2009). Liver-specific loss of lipolysis-stimulated lipoprotein receptor triggers systemic hyperlipidemia in mice. Diabetes 58, 1040–1049.

Papatheodorou, P., Carette, J.E., Bell, G.W., Schwan, C., Guttenberg, G., Brummelkamp, T.R., and Aktories, K. (2011). Lipolysis-stimulated lipoprotein receptor (LSR) is the host receptor for the binary toxin Clostridium difficile transferase (CDT). Proceedings of the National Academy of Sciences of the United States of America 108, 16422–16427.

Reaves, D.K., Hoadley, K.A., Fagan-Solis, K.D., Jima, D.D., Bereman, M., Thorpe, L., Hicks, J., McDonald, D., Troester, M.A., Perou, C.M., et al. (2017). Nuclear Localized LSR: A Novel Regulator of Breast Cancer Behavior and Tumorigenesis. Mol Cancer Res 15, 165–178.

Resnik-Docampo, M., Koehler, C.L., Clark, R.I., Schinaman, J.M., Sauer, V., Wong, D.M., Lewis, S., D’Alterio, C., Walker, D.W., and Jones, D.L. (2017). Tricellular junctions regulate intestinal stem cell behaviour to maintain homeostasis. Nature cell biology 19, 52–59.

Sato, T., Vries, R.G., Snippert, H.J., van de Wetering, M., Barker, N., Stange, D.E., van Es, J.H., Abo, A., Kujala, P., Peters, P.J., et al. (2009). Single Lgr5 stem cells build crypt-villus structures in vitro without a mesenchymal niche. Nature 459, 262–265.

Schlegelmilch, K., Mohseni, M., Kirak, O., Pruszak, J., Rodriguez, J.R., Zhou, D., Kreger, B.T., Vasioukhin, V., Avruch, J., Brummelkamp, T.R., et al. (2011). Yap1 acts downstream of alpha-catenin to control epidermal proliferation. Cell 144, 782–795.

Serra, D., Mayr, U., Boni, A., Lukonin, I., Rempfler, M., Challet Meylan, L., Stadler, M.B., Strnad, P., Papasaikas, P., Vischi, D., et al. (2019). Self-organization and symmetry breaking in intestinal organoid development. Nature 569, 66–72.

Shimada, H., Abe, S., Kohno, T., Satohisa, S., Konno, T., Takahashi, S., Hatakeyama, T., Arimoto, C., Kakuki, T., Kaneko, Y., et al. (2017). Loss of tricellular tight junction protein LSR promotes cell invasion and migration via upregulation of TEAD1/AREG in human endometrial cancer. Sci Rep 7, 37049.

Sohet, F., Lin, C., Munji, R.N., Lee, S.Y., Ruderisch, N., Soung, A., Arnold, T.D., Derugin, N., Vexler, Z.S., Yen, F.T., et al. (2015). LSR/angulin-1 is a tricellular tight junction protein involved in blood-brain barrier formation. The Journal of cell biology 208, 703–711.

Sugawara, T., Furuse, K., Otani, T., Wakayama, T., and Furuse, M. (2021). Angulin-1 seals tricellular contacts independently of tricellulin and claudins. The Journal of cell biology 220.

Takahashi, Y., Serada, S., Ohkawara, T., Fujimoto, M., Hiramatsu, K., Ueda, Y., Kimura, T., Takemori, H., and Naka, T. (2021). LSR promotes epithelial ovarian cancer cell survival under energy stress through the LKB1-AMPK pathway. Biochem Bioph Res Co 537, 93–99.

Totaro, A., Panciera, T., and Piccolo, S. (2018). YAP/TAZ upstream signals and downstream responses. Nature cell biology 20, 888–899.

Underwood, M.A. (2012). Paneth cells and necrotizing enterocolitis. Gut microbes 3, 562–565.

Wehkamp, J., and Stange, E.F. (2020). An Update Review on the Paneth Cell as Key to Ileal Crohn’s Disease. Frontiers in immunology 11, 646.

Yen, F.T., Masson, M., Clossais-Besnard, N., Andre, P., Grosset, J.M., Bougueleret, L., Dumas, J.B., Guerassimenko, O., and Bihain, B.E. (1999). Molecular cloning of a lipolysis-stimulated remnant receptor expressed in the liver. The Journal of biological chemistry 274, 13390–13398.

Zhao, B., Li, L., Tumaneng, K., Wang, C.Y., and Guan, K.L. (2010). A coordinated phosphorylation by Lats and CK1 regulates YAP stability through SCF(beta-TRCP). Genes & development 24, 72–85.

Zhou, D., Zhang, Y., Wu, H., Barry, E., Yin, Y., Lawrence, E., Dawson, D., Willis, J.E., Markowitz, S.D., Camargo, F.D., et al. (2011). Mst1 and Mst2 protein kinases restrain intestinal stem cell proliferation and colonic tumorigenesis by inhibition of Yes-associated protein (Yap) overabundance. Proceedings of the National Academy of Sciences of the United States of America 108, E1312–1320.

