## Supplementary Material and Methods for "LSR Targets YAP to Modulate Intestinal Paneth Cell Differentiation"

### Animals

All animals were housed in micro-isolator cages and received food and water freely. The laboratory temperature was 24±1°C, and the relative humidity was 40-60%. The protocol in this study was approved by the Committee on the Ethics of Animal Care and Use of Binzhou Medical University. The study was conducted following the Guide for the Care and Use of Animals in Research of the People's Republic of China. All animal studies were performed under isoflurane anesthesia, and every effort was made to minimize suffering. C57BL/6J mice were obtained from Vital River Laboratory (China).

For the generation of $Lsr^{loxP/loxP}$ mice, targeting vector was constructed by inserting one SDA (self-deletion anchor)-flanked neomycin cassette and two loxP sites flanking exons 1~2 of Lsr and then electroporating into embryonic stem cells from C57BL/6J mice (conducted by Cyagen Biosciences Inc, China). With one subsequent cross with C57BL/6J animals, the Neo transgene was removed and the obtained $Lsr^{loxP/-}$ mice were then intercrossed in order to generate the $Lsr^{loxP/loxP}$ mice. $Lsr^{loxP/loxP}$ mice were crossed with villin-Cre mice (Jackson Laboratory, 021504) [1], a mouse Cre line that drives Cre recombination throughout the intestinal epithelium from embryonic day 12.5, to remove Lsr from intestinal epithelium ($Lsr^{loxP/loxP}$/villin-$Cre^{+/-}$, $Lsr^{vill\ KO}$). Homozygous with villin-Cre negative litter mates served as controls. $Lsr^{loxP/loxP}$ mice were crossed with *CAG-CreER* mice (Jackson Laboratory, 004682) [2] to generate systemic *Lsr* knockout mice $Lsr^{loxP/loxP}/CAG\text{-}Cre^{+/-}$ ($Lsr^{-/-}$), homozygous with *CAG-Cre* negative litter mates served as controls. $Lsr^{loxP/loxP}$ mice were crossed with *Lgr5-CreERT2* mice (GemPharmatech, T003768) to generate ISC-specific LSR knockout mice ($Lsr^{loxP/loxP}$/Lgr5-$CreERT2^{+/-}$, $Lsr^{ISC\ KO}$), homozygous with *Lgr5-CreERT2* negative litter mates served as controls. To induce the knockout of genes, tamoxifen suspended in corn oil was administered intraperitoneally at a concentration of 10 mg/ml and dosed at 100mg/kg once every 24 hours for a total of 5 consecutive days. 15 days post last tamoxifen injection, tissues were harvested for qRT-PCR, Western blot, H&E or immunofluorescence staining analysis.

### Patient sample

Human intestinal tissues were collected after approval by the Binzhou Medical University Review Board. De-identified formalin-fixed paraffin-embedded tissue sections of NEC were compared with healthy tissue margins resected for indications other than NEC (intestinal obstruction or spontaneous intestinal perforation (n = 10). Written informed consent for research was obtained from donors before tissue acquisition. The patients were from both sexes, between 4 and 49 days of born, and weighed between 0.9 and 1.15 kg.

**Cell lines, plasmids, and antibodies**

Human HEK293 cells were obtained from ATCC (USA) and iPSCs were obtained from National Collection of Authenticated Cell Cultures (China), and cultured according to the distributor's recommendations. The expression plasmids containing mouse Lsr (pCMV6-LSR, NM_001164185, MR225038), human LSR (pCMV6-LSR, NM_NM_205834, RC207960), mouse Yap (pCMV6-YAP, NM_001171147, MR226049), mouse 14-3-3 zeta (pCMV6-14-3-3 zeta, NM_011740, MR203074), mouse 14-3-3gamma (pCMV6-14-3-3 gamma, NM_018871, MR203103), mouse Ppp1ca (pCMV6-Ppp1ca, NM_031868, MR204856), mouse PP2AC (pCMV6- Pp2ac, NM_019411, MR204384) genes, and the knockdown plasmids containing YAP mouse shRNA plasmid (pGFP-C-shLenti, 22601) were purchased from Origene Technologies (USA). The antibodies used in this study are summarized in Supplemental Table 1.

**Intestinal permeability analysis**

Intestinal permeability was evaluated with fluorescein isothiocyanate (FITC)-Dextran 4000 (MCE, HY-128868A, China) test. Briefly, mice were fasted for 6 h and then FITC-dextran 4000 (440 mg/kg body wt) was administered by gavage. Blood was collected through tail vein 0h, 0.5h, 1h, 2h, 4h and 8h after gavage, and serum was isolated. The analysis for the fluorescence intensity of FITC-dextran 4000 in plasm was carried out with a multimode plate reader (PerkinElmer EnSpire Plate Reader, Waltham, MA, USA) (excitation, 485 nm; emission, 520 nm). Standard curves for calculating the FITC-dextran concentration were obtained by diluting FITC-dextran in PBS.

**TNBS-induced NEC**

2, 4, 6-Trinitrobenzene sulfonic acid (TNBS) were used to develop NEC [3]. Briefly, TNBS was injected into 14-day-old C57BL/6J mice (n = 8) (50 mg/kg total body weight, dissolved in 30% weight/volume ethanol by gavage and enema). Animals were monitored every 3h and were euthanized if they developed signs of disease or at 48h using $CO_2$ inhalation followed by cervical dislocation. Small intestine was harvested and processed for histological analyses. As previously described [4], intestinal injury was confirmed by histopathological grading and was assigned by a blinded reviewer: grade 0 = no injury; grade 1 = mild injury: disruption of villus tips or mild separation of lamina propria in the small intestine; grade 2 = moderate injury: villus disruption, clear separation of lamina propria and/or edema in the submucosa; grade 3 = severe injury: transmural injury in the small intestine. Others were collected for qRT-PCR and immunofluorescence analysis. Control animals (n=8) received vehicle alone (30% ethanol).

***In vivo* administration of recombinant AAV**

AAVrh10 has been proved to be the most efficient rAAV serotype for intestine gene delivery [5]. 7-day-old C57BL/6J mice received with $1 \times 10^8$ MOI virus genomic particles of Rh10-pHBAAV-CMV-MCS-3flag-T2A-ZsGreen (AAV-CTL) or Rh10-pHBAAV-CMV-Lsr-3flag-T2A-ZsGreen (AAV-*Lsr*) (prepared by Hanbio Biotechnology Inc, China) by intraperitoneal injection (n = 8). 1 week after transduction, mice were treated with TNBS to construct NEC models as described above.

To silence YAP *in vivo*, 1-week-old $Lsr^{vill\ KO}$ mice were injected intraperitoneally with a single dose of $2 \times 10^8$ MOI particles of lentivirus/pHBLV-U6-MCS-CMV-ZsGreen-PGK-PURO (CTL) or lentivirus/pHBLV-U6-YAP-siRNA-CMV-ZsGreen-PGK-PURO (*Yap* shRNA) (prepared by Hanbio Biotechnology Inc, China) (n = 8). Mice were monitored every day for general wellbeing. 2 weeks after transduction, mice were euthanized using $CO_2$ inhalation followed by cervical dislocation. Small intestine tissues were harvested for RNA isolation and immunofluorescence and histological staining.

***In vivo* administration of YAP inhibitor verteporfin**

1-week-old $Lsr^{vill\ KO}$ mice were injected intraperitoneally with verteporfin (100 mg/kg body weight) every 2 days for 2 weeks, (n = 8). Mice were monitored every day for general wellbeing. After 2 weeks, mice were euthanized using $CO_2$ inhalation followed by cervical dislocation. Small intestine tissues were harvested for RNA isolation and immunofluorescence and histological staining.

**Histologic analysis**

Human and mouse intestine tissues were fixed overnight at 4°C in 4% paraformaldehyde and embedded in paraffin. Intestine sections (5 μm thickness) were prepared by a routine procedure, then standard histological staining with H&E and periodic acid-Schiff (PAS). Transmission electron microscopy (TEM) was performed on glutaraldehyde-fixed, epoxy-embedded intestine samples and stained with uranyl acetate and lead citrate.

**Immunofluorescence staining**

The small intestine (duodenal, jejunal and ileal) cross sections were embedded in OCT. Small intestine cryosections (8 μm) were mounted onto glass slides, fixed in cold methanol or acetone for 20 min at -20°C. For immunofluorescence of mouse or human organoid, organoid on collagen I coated confocal dish were fixed with 4% paraformaldehyde. And then washed three times with PBS for 10 min each time, and blocked for 1 h at room temperature in 10% BSA diluted in PBS (containing 0.01% Tween 20). Then incubated overnight at 4°C with appropriate primary antibodies. Washed as before and incubated using FITC and/or rhodamine (TRITC) conjugated secondary antibodies (Invitrogen), followed by DAPI (Sigma) nuclear staining. And then sealed with ProLong$^{TM}$ Gold antifade reagent and analyzed using a confocal laser microscope (LSM 880 Basic Operation, Germany).

For human Paraffin section (5 μm thickness), immunostaining was preceded by antigen retrieval in EDTA buffer (pH 8.0), and blocked for 1 h at room temperature in 10% BSA. The primary antibodies were anti-human: LSR, MPTX2, OFLM4 and incubated overnight at 4°C. Then incubated with FITC and/or rhodamine (TRITC) conjugated secondary antibodies (Invitrogen) for 1.5 h at room temperature, followed by DAPI (Sigma) nuclear staining. The images were acquired using confocal laser microscope (LSM 880 Basic Operation, Germany).

**RNA-sequencing (RNA-seq)**

Total RNA was extracted using Trizol reagent (Invitrogen, 15596018) from intestine of $Lsr^{ctrl}$ and $Lsr^{vill}$

$^{KO}$ mice at 4 weeks after birth. RNA-seq was performed on an Illumina Novaseq platform by Novogene (China). Paired-end clean reads were aligned to the mouse reference genome (Ensemble_GRCm38.90) with TopHat (version 2.0.12), and the aligned reads were used to quantify mRNA expression by using HTSeq-count (version 0.6.1). Differential expression analysis of two groups was performed using the DESeq2 R package (1.16.1). The resulting P-values were adjusted using the Benjamini and Hochberg's approach for controlling the false discovery rate. Genes with an adjusted *P-value <0.05* found by DESeq2 were assigned as differentially expressed.

**Crypt and Lgr5$^+$ ISCs isolation and culture**

Small intestines crypt cultures were prepared using previously described methods with modifications [6,7]. In short, small intestines were first flushed with PBS, then cut longitudinally. The tissue was chopped into 2-3 mm pieces, and further washed with cold PBS. The tissue fragments were incubated in 5 mM EDTA-PBS solution on ice. Then EDTA was removed, the supernatant fractions enriched for crypts were passed through a 70-micron strainer (BD Biosciences) to remove the remaining cells of the intestinal villi fraction. The filtered fractions were then centrifuged to collect the crypt fraction. Crypts isolated from the small intestine of Lsr$^{vill\ KO}$ and Lsr$^{ctrl}$ mice was resuspended into matrigel with DMEM/F12 medium (1:1 ratio) and plated into 24 well plates [8]. Crypt fraction organoids were kept in IntestiCult Organoid Growth Medium (STEMCELL Technologies) with 100 μg/ml Penicillin-Streptomycin for amplification and maintenance.

For ISCs isolation, crypts were isolated from the small intestine of Lsr$^{vill\ KO}$ and Lsr$^{ctrl}$ mice as described previously in the methods. ISCs were isolated from the small intestine by immunomagnetic beads. In short, the crypt suspensions were centrifuged and resuspended in undiluted TrypLE Express (Invitrogen) supplemented with DNase I (2000 U/μl, Roche), and 10 μM Y27632 (MCE) and incubated at 37°C for 8min. Then, cold DMEM/F12 (STEMCELL Technologies) were added to the samples and triturated twice, followed by centrifugated and resuspend the cell pellet in DMEM/F12. Positive selection of ISCs were achieved using magnetic beads (1 μm diameter, Invitrogen) coated with a mouse monoclonal antibody against LGR5 (invitrogen). For PA treatment, Lgr5$^+$ ISCs isolated from mouse small intestine were incubated in serum-free medium supplemented with 0.3 mM PA for 24 hours. And then cells were harvested for RT-PCR, Western blot or immunofluorescence staining analysis.

**Lentivirus transduction of organoids**

For Lentivirus-Green-PURO-YAP (YAP-KD) virus infection experiments, cells recovery solution (Corning) was used to harvest organoids and mechanically dissociated in Basal Media using a 10 ml Pipette for 5-7 times up and down. Then, centrifuged the organoids at 250 × g for 5 min, 4°C, and resuspend use 400 μl full growth medium containing 1 × $10^7$ MOI YAP-KD Lentivirus, then add to 100 μl matrigel together. The mixture was incubated at 37°C for 5 h, mixing every 30 min. Later, the mixture was centrifuged at 250 × g for 5 min at 4°C and organoids were seeded in matrigel and cultured for 3 days before harvesting for RNA isolation and immunofluorescence staining.

**shRNA molecular cloning, screening and retrovirus production**

The shRNA hairpin oligonucleotides, complementary to all the variants of human Lsr mRNA sequences (NM_015925.7, NM_205834.4, NM_205835.4, NM_001260489.2, NM_001260490.2, and NM_001385215.1) (Table S2), were synthesized by Sangon Biotechnology Inc, annealed, and cloned into the moloney murine leukemia retrovirus vector pSIREN (Clontech), downstream of the human small nuclear ribonucleoprotein U6 promoter to create the *LSR* shRNA constructs. Vesicular stomatitis virus glycoprotein (VSV-G) pseudo typed retroviruses were produced in HEK293 cells and used to infect target cells at a titer of 1 × $10^7$ CFU/ml, as described previously [9].

**Maintenance of iPSCs and *LSR* knockdown**

Human iPSCs were maintained on matrigel (Corning) in mTesR1 medium (STEMCELL Technologies). Cells were passaged approximately every 4-5 days. To passage iPSCs, they were washed with serum free DMEM/F12 medium (STEMCELL Technologies) and incubated in DMEM/F12 with 1 mg/ml dispase (Invitrogen) until cell edges started to detach from the dish. And then replaced with mTesR1. pipette gently up and down to make the cells into small clumps and passaged onto fresh matrigel-coated plates. For virus transfection experiments, above cells add full growth medium containing 1 × $10^7$ MOI above virus, the cells were incubated at 37°C for 12 h. Then, the medium was changed with fresh complete medium and cultured for 48 h before harvesting for RNA isolation and organoid culture.

**Differentiation of iPSCs into human intestine organoid**

Differentiation of human iPSCs into organoid was run in accordance with the manufacturer's instructions (STEMCELL Technologies). In brief, $1.5 \times 10^5$ iPSCs per ml were plated on matrigel coated tissue culture plate (Thermo Fisher). Medium was changed to STEMdiff™ Endoderm Basal Medium (STEMCELL Technologies) containing STEMdiff™ Definitive Endoderm Supplement CJ (STEMCELL Technologies). On day 3, medium was changed to STEMdiff™. Endoderm Basal Medium containing STEMdiff™. Gastrointestinal Supplement PK + STEMdiff™. Gastrointestinal Supplement UB (STEMCELL Technologies). On day 4-9, medium was changed to Intestinal OGM (STEMCELL Technologies) and incubate at 37°C with 5% $CO_2$ and 95% humidity.

**Quantitative real time PCR**

The small intestine samples were homogenized in Trizol (Ambion, USA) to get total RNA. RNA was reverse transcribed by using a reverse transcription system kit according to the instructions of the manufacturer (Takara, Japan) followed by quantitative real time PCR (qRT-PCR) amplification using SYBR Green PCR Master Mix (ABI, USA) and the Thermo Fisher QuantStudio3 system. The Supplemental Table 2 contains the primer sequences used in this study. The expression levels of each mRNA were calculated after normalizing to those of β-actin. Results were expressed as $2^{-\Delta Ct}$ values with $\Delta CT = Ct_{gene} - Ct_{\beta\text{-actin}}$.

**Western blot analysis**

Total protein extracts were obtained by lysing isolated ISCs or small intestine tissues in RIPA lysis buffer (Sangon Biotech) containing PMSF and phosphatase inhibitors (Sangon Biotech) on ice, supernatants were collected after centrifugation at 4°C for 10 min at 12,000 × g. Then, the protein concentration was assessed using a BCA protein assay (Takara). Proteins (50 µg) were separated in 10% SDS-PAGE and then transferred onto PVDF membranes (0.22 µm), and then blocked with 5% skim milk dissolved in TBST and incubated with the primary antibody at 4°C overnight. HRP-conjugated secondary antibodies (Thermo Fisher) followed by ECL (Thermo Fisher) incubation allowed protein band detection. The integration of all blots images was performed on Adobe Illustrator 2021. Image J v1.8.0 was used to quantify western blotting results.

**Co-Immunoprecipitation (Co-IP)**

Isolated ISCs or HEK293 cells transfected with expressing vectors for LSR, YAP, and 14-3-3 zeta; LSR, YAP, and 14-3-3 gamma; LSR, YAP, and PPP1CA; LSR, YAP, and PP2Ac; and YAP, 14-3-3 zeta and PP2Ac were lysed in 50mM Tris (pH 8.0) by 25–30 repeated passages through a 25-gauge needle, followed by centrifugation at 5,000 × g for 20 min, 4°C. The membranes of lysis were extracted using CSK buffer (1% Triton X-100; 150 mM NaCl; 50 mM Tris, pH 8.0; and protease inhibitors). The membrane extract was precleared by incubation with protein A/G-sepharose (invitrogen) prior to Co-IP. The precleared membrane extract was incubated for 16 h at 4°C with anti-LSR, anti-YAP, anti-14-3-3 zeta, anti-14-3-3 gamma, anti-PPP1CA, anti-PP2Ac, anti-mouse IgG, or anti-Rabbit IgG antibodies. Antibody-bound material was pelleted with protein A/G-sepharose, washed three times with CSK buffer, and detected by immunoblotting.

**LC-MS/MS Analysis**

Follow the above Co-IP method, isolated ISCs and extracted the membranes of lysis. The precleared membrane extract was incubated for 16 h at 4°C with anti-LSR and anti-mouse IgG (as negative control) antibodies. Antibody-bound material was pelleted with protein A/G-sepharose, washed three times with CSK buffer, and performing SDS-PAGE, and then cut the gel and detected.

The separated peptides were analyzed by Q Exactive HF-X mass spectrometer (Thermo Fisher) by Novogene (China). The resulting spectra from each fraction were searched separately against uniprot database by the search engines: Proteome Discoverer 2.2 (PD 2.2, Thermo). The search parameters are set as follows: mass tolerance for precursor ion was 10 ppm and mass tolerance for product ion was 0.02 Da. Carbamidomethyl was specified in PD 2.2 as fixed modifications. Oxidation of methionine (M) and acetylation of the N-terminus were specified in PD 2.2 as variable modifications. A maximum of 2 missed cleavage sites were allowed. The identified protein contains at least 1 unique peptide with FDR no more than 1%. Proteins containing similar peptides that could not be distinguished by MS/MS analysis were identified as a same protein group.

**Data availability**

The raw RNA-seq data were deposited in NCBI sequence read archive (SRA) database, with the accession number PRJNA769233. Additional details on protocols and databases/datasets used in this study are available from the corresponding authors on reasonable request.

**Statistical analysis**

The significance of differences between groups was tested by Prism 8 (GraphPad Software Inc.). Statistical analysis was performed using the unpaired *t* test to determine differences between two groups and ANOVA to compare data among groups. P-values less than 0.05 were interpreted as statistically significant. All data are presented as mean ± SEM and other details such as the number of replicates and the level of significance is mentioned in figure legends and supplementary information.

**Reference**


1 Madison, B. B. *et al.* Cis elements of the villin gene control expression in restricted domains of the vertical (crypt) and horizontal (duodenum, cecum) axes of the intestine. *The Journal of biological chemistry* **277**, 33275-33283, doi:10.1074/jbc.M204935200 (2002).

2 Hayashi, S. & McMahon, A. P. Efficient recombination in diverse tissues by a tamoxifen-inducible form of Cre: a tool for temporally regulated gene activation/inactivation in the mouse. *Developmental biology* **244**, 305-318, doi:10.1006/dbio.2002.0597 (2002).

3 MohanKumar, K. *et al.* Gut mucosal injury in neonates is marked by macrophage infiltration in contrast to pleomorphic infiltrates in adult: evidence from an animal model. *American journal of physiology. Gastrointestinal and liver physiology* **303**, G93-102, doi:10.1152/ajpgi.00016.2012 (2012).

4 Namachivayam, K. *et al.* Targeted inhibition of thrombin attenuates murine neonatal necrotizing enterocolitis. *Proceedings of the National Academy of Sciences of the United States of America* **117**, 10958-10969, doi:10.1073/pnas.1912357117 (2020).

5 Polyak, S. *et al.* Identification of adeno-associated viral vectors suitable for intestinal gene delivery and modulation of experimental colitis. *American journal of physiology. Gastrointestinal and liver physiology* **302**, G296-308, doi:10.1152/ajpgi.00562.2010 (2012).

6 Goga, A. *et al.* miR-802 regulates Paneth cell function and enterocyte differentiation in the mouse small intestine. *Nature communications* **12**, 3339, doi:10.1038/s41467-021-23298-3 (2021).

7 Gjorevski, N. *et al.* Designer matrices for intestinal stem cell and organoid culture. *Nature* **539**, 560-564, doi:10.1038/nature20168 (2016).

8 Serra, D. *et al.* Self-organization and symmetry breaking in intestinal organoid development. *Nature* **569**, 66-72, doi:10.1038/s41586-019-1146-y (2019).

9 Gong, Y. *et al.* The Cap1-claudin-4 regulatory pathway is important for renal chloride reabsorption and blood pressure regulation. *Proceedings of the National Academy of*

*Sciences of the United States of America* **111**, E3766-3774, doi:10.1073/pnas.1406741111 (2014).
