## Supplementary Figures for "LSR Targets YAP to Modulate Intestinal Paneth Cell Differentiation"

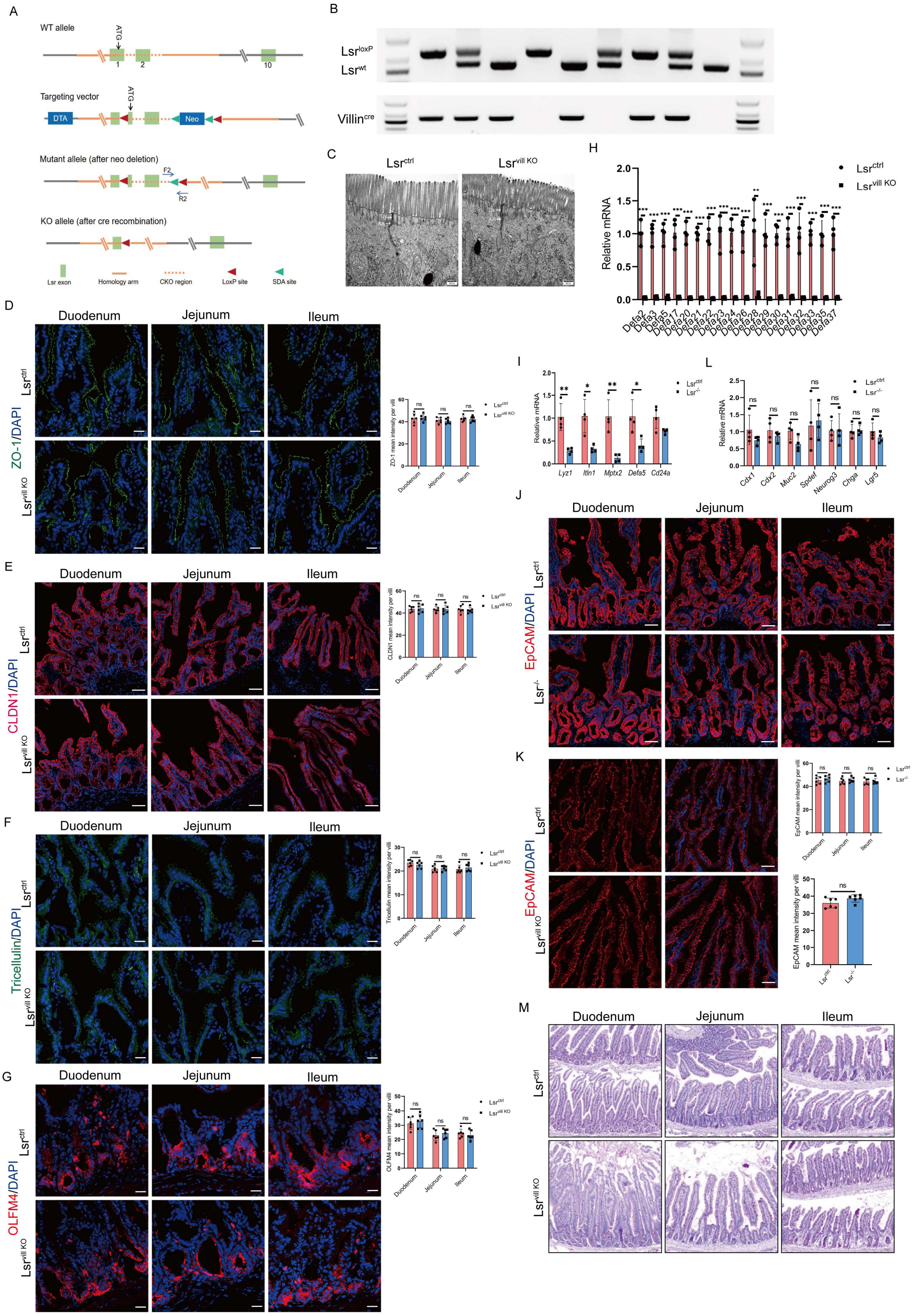


**Supplementary Figure 1. LSR ablation results in increased numbers of proliferating cells and loss of Paneth cell lineage in the small intestine.**

(A) Gene targeting strategy. The diagram showing the wild-type Lsr locus, the targeting construct, the mutant allele after neo deletion, and the knockout (KO) allele after Cre recombination. In the targeting vector, the neo cassette was flanked by SDA (self-deletion anchor) sites, and exons 1~2 of *Lsr* was flanked by loxP sites. The first loxP site was placed upstream of the ATG start codon with 8 bp. Diphtheria toxin A (DTA) was used for negative selection. (B) PCR verification of offspring tails, demonstrating WT (438 bp), heterozygous (438 bp and 551 bp), and homozygous (551 bp) alleles. The primer F2 is located on cKO (conditional knock out) region. The primer R2 is located downstream of the second loxP site. The villin Cre recombinase transgene was identified as a 430 bp PCR product. (C) TEM in small intestines from Lsr$^{\mathrm{ctrl}}$ and Lsr$^{\mathrm{vill\ KO}}$ mice. Arrows indicated tight junction (TJ) and adherens junction (AJ). (D-F) ZO-1 (D), CLDN1 (E), and tricellulin (F) immunofluorescence and quantification analysis in small intestines from Lsr$^{\mathrm{ctrl}}$ and Lsr$^{\mathrm{vill\ KO}}$ mice (n=6). (G) OLFM4 immunofluorescence and quantification analysis in duodenal, jejunal and ileal segments from Lsr$^{\mathrm{ctrl}}$ and Lsr$^{\mathrm{vill\ KO}}$ mice (n=6). (H) Expression of *Defa* family mRNA assessed by qRT–PCR in small intestines from Lsr$^{\mathrm{ctrl}}$ and Lsr$^{\mathrm{vill\ KO}}$ mice (n=4). (I) Expression of *Lyz1, Itln1, Mptx2, Defa5* and *Cd24a* mRNA assessed by qRT–PCR in small intestines from Lsr$^{\mathrm{ctrl}}$ and Lsr$^{-/-}$ mice (n=4). (J and K) EpCAM immunofluorescence and quantification analysis in small intestines from Lsr$^{\mathrm{ctrl}}$ and Lsr$^{-/-}$mice (J) and Lsr$^{\mathrm{ctrl}}$ and Lsr$^{\mathrm{vill\ KO}}$ mice (K) (n=6). (L) qRT–PCR analysis the mRNA expression of columnar absorptive cell markers (*Cdx1* and *Cdx2*), goblet cell markers (*Muc2* and *Spdef*), enteroendocrine cell markers (*Neurog3* and *Chga*), and ISCs marker (*Lgr5*) in small intestines from Lsr$^{\mathrm{ctrl}}$ and Lsr$^{-/-}$ mice (n=4). (M) Representative PAS-stained small intestine from Lsr$^{\mathrm{ctrl}}$ and Lsr$^{\mathrm{vill\ KO}}$ mice. Scale bars: C, 500 nm; E, F and G, 20 μm; J and K, 50 μm; M, 120 μm. *$P < 0.05$, **$P < 0.01$, ***$P < 0.001$.

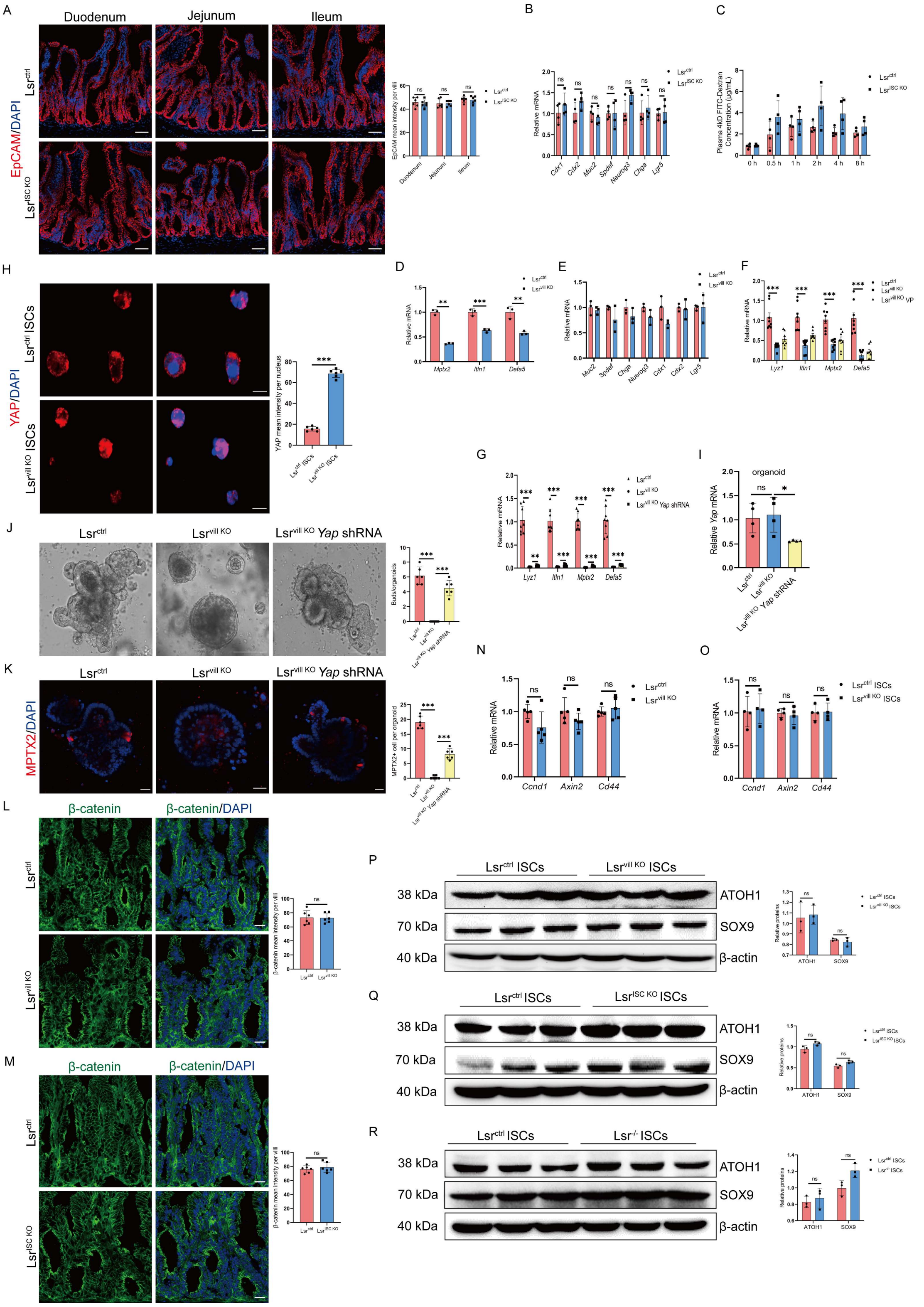


**Supplementary Figure 2. LSR regulated Lgr5$^+$ ISCs differentiated into Paneth cell via the activation of YAP.**

(A) EpCAM immunofluorescence and quantification analysis in small intestines from the Lsr$^{ctrl}$ and Lsr$^{ISC\ KO}$ mice (n=6). (B) qRT–PCR analysis the mRNA expression of columnar absorptive cell markers

(*Cdx1* and *Cdx2*), goblet cell markers (*Muc2* and *Spdef*), enteroendocrine cell markers (*Neurog3* and *Chga*), and ISCs marker (*Lgr5*) in small intestines from Lsr$^{ctrl}$ and Lsr$^{ISC\ KO}$ mice (n=4). (C) Intestinal permeability to 4 kDa FITC-dextran (FD4) of Lsr$^{ctrl}$ and Lsr$^{ISC\ KO}$ mice (n=4). (D and E) qRT–PCR analysis of the mRNA expression of Paneth cell markers (*Mptx2, Itln1,* and *Defa5*) (D) and other IEC markers including columnar absorptive cell markers (*Cdx1* and *Cdx2*), goblet cell markers (*Muc2* and *Spdef*), enteroendocrine cell markers (*Neurog3* and *Chga*), and ISCs marker (*Lgr5*) (E) in organoids established from intestinal crypts of Lsr$^{ctrl}$ and Lsr$^{vill\ KO}$ mice (n=3). (F and G) qRT–PCR analysis of the mRNA expression of Paneth cell markers in small intestines from Lsr$^{ctrl}$ mice, Lsr$^{vill\ KO}$ mice and Lsr$^{vill\ KO}$ mice treated with VP (F) and Lsr$^{ctrl}$ mice, Lsr$^{vill\ KO}$ mice and Lsr$^{vill\ KO}$ mice treated with lentivirus expressing *Yap* shRNA (G) (n=8). (H) YAP immunofluorescence and quantification analysis in ISCs isolated from Lsr$^{ctrl}$ and Lsr$^{vill\ KO}$ mice (n=6). (I) qRT–PCR analysis for *Yap* mRNA in crypt organoids generated from Lsr$^{ctrl}$ mice, Lsr$^{vill\ KO}$ mice, and Lsr$^{vill\ KO}$ mice organoids treated with lentivirus expressing *Yap* shRNA (n=4). (J) Bright field images and quantification analysis of Lsr$^{ctrl}$ crypt organoids, Lsr$^{vill\ KO}$ crypt organoids with and without transduction with lentivirus expressing *Yap* shRNA (n=6). (K) MPTX2 immunofluorescence and quantification analysis of Lsr$^{ctrl}$ crypt organoids, Lsr$^{vill\ KO}$ crypt organoids with and without transduction with lentivirus expressing *Yap* shRNA (n=6). (L and M) β-catenin immunofluorescence and quantification analysis in small intestines from Lsr$^{ctrl}$ and Lsr$^{vill\ KO}$ mice (L) and Lsr$^{ctrl}$ and Lsr$^{ISC\ KO}$ mice (M) (n=6). (N and O) mRNA expression of *Ccnd1, Axin2* and *Cd44* in small intestines (N) and ISCs (O) from Lsr$^{ctrl}$ and Lsr$^{vill\ KO}$ mice (n=4). (P-R) Western blot and quantification analysis of ATOH1 and SOX9 protein expression in isolated ISCs from Lsr$^{ctrl}$ and Lsr$^{vill\ KO}$ mice (P), Lsr$^{ctrl}$ and Lsr$^{ISC\ KO}$ mice (Q), and Lsr$^{ctrl}$ and Lsr$^{-/-}$ mice (R) (n=3). Scale bars: F, 50 μm; H, 10μm; J, 125 μm; K, L, M, 20 μm. *$P < 0.05$, **$P < 0.01$, ***$P < 0.001$.

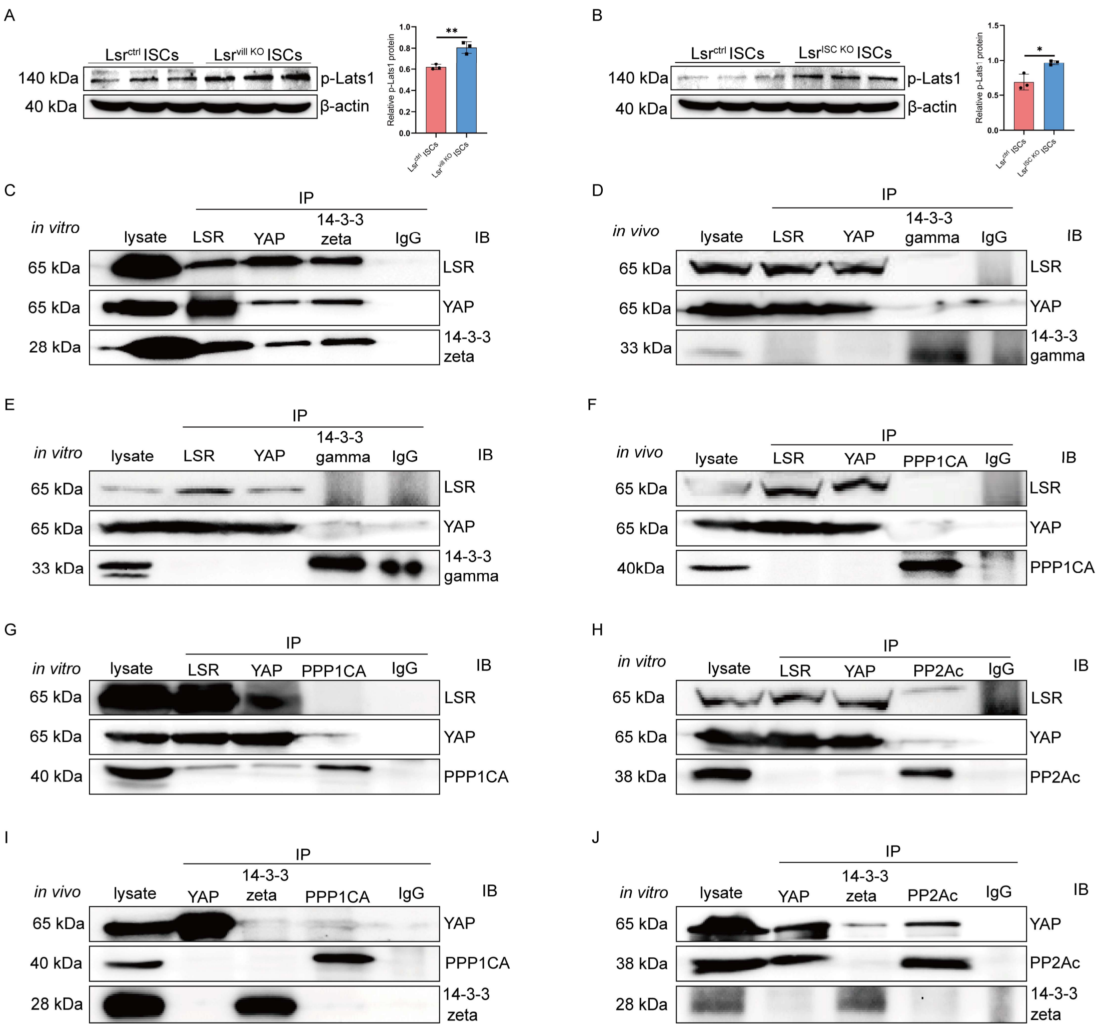


**Supplementary Figure 3. 14-3-3 zeta and PP2Ac are involved in the regulation of YAP by LSR.** (A and B) Western blot and quantification analysis for p-Lats1 in isolated ISCs from $Lsr^{ctrl}$ and $Lsr^{vill\ KO}$ mice (A) and $Lsr^{ctrl}$ and $Lsr^{ISC\ KO}$ mice (B) (n=3). (C) Co-IP showing that LSR, YAP and 14-3-3 zeta interact with each other in multiply transfected HEK293. (D) Co-IP showing that endogenous YAP interacts with endogenous LSR, but not 14-3-3 gamma in isolated ISCs from $Lsr^{ctrl}$ mice. (E) Co-IP showing that YAP interacts with LSR, but not 14-3-3 gamma in multiply transfected HEK293. (F) Co-IP showing that endogenous YAP interacts with endogenous LSR, but not PPP1CA in isolated ISCs from $Lsr^{ctrl}$ mice. (G) Co-IP showing that YAP interacts with LSR, but not PPP1CA in multiply transfected HEK293. (H) Co-IP showing that YAP interacts with LSR, but not PP2Ac in multiply transfected HEK293. (I) Co-IP showing that endogenous YAP, PPP1CA, and 14-3-3 zeta couldn't interact with each other in isolated ISCs from $Lsr^{vill\ KO}$ mice. (J) Co-IP showing that YAP interacts with PP2Ac, but not 14-3-3 zeta in multiply transfected HEK293 transduced with lentivirus expressing *Lsr* shRNA. In all of the Co-IP, the antibodies used for immunoprecipitation are shown above the lanes; the antibody for immunoreaction visualization are shown on the right. $^{*}P < 0.05$, $^{**}P < 0.01$, $^{***}P < 0.001$.

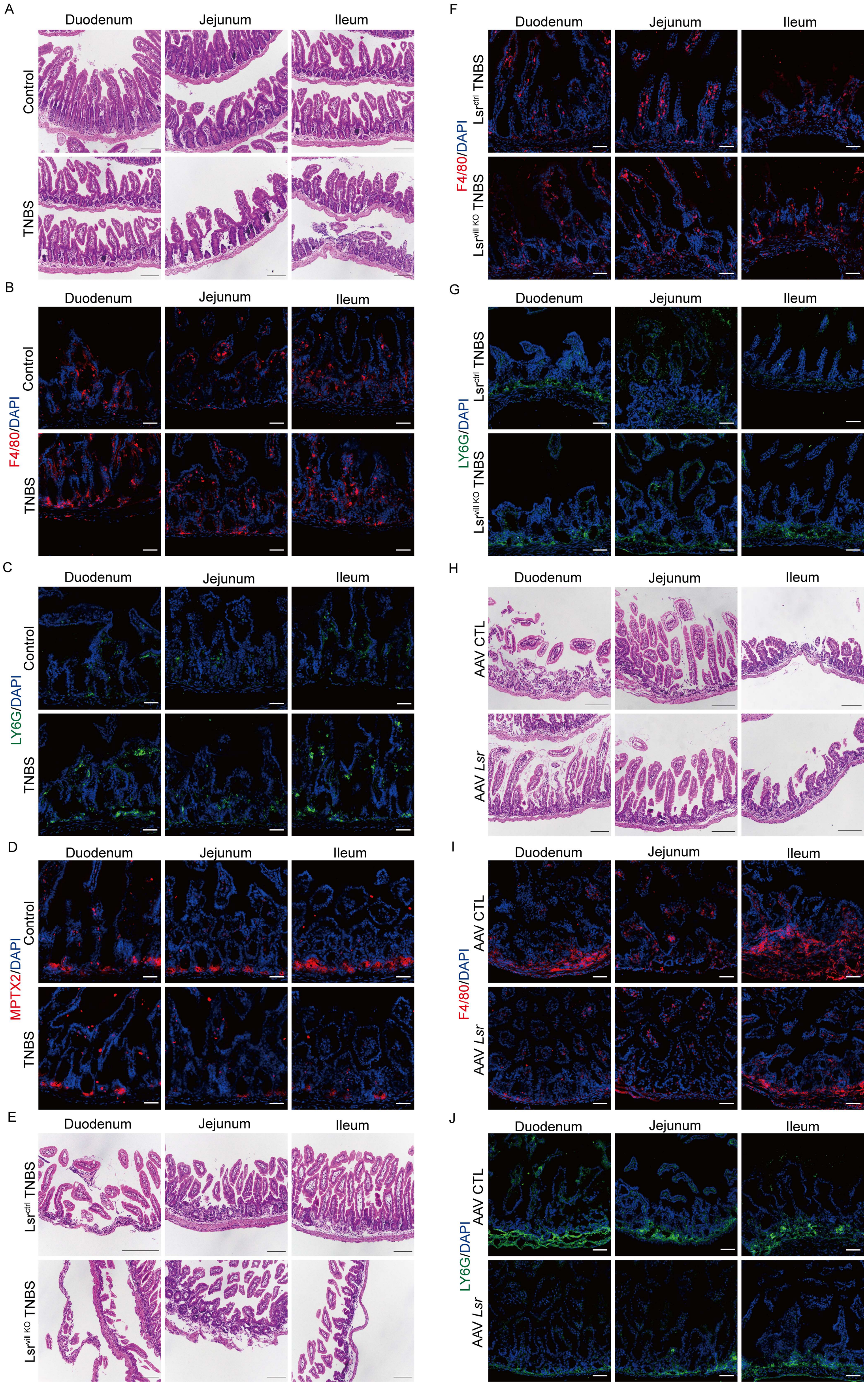


**Supplementary Figure 4. Loss of Lsr exacerbates inflammation in a mouse model of NEC.**

(A) Representative H&E-stained small intestine from control and TNBS-treated mice. (B and C) F4/80 (B) and LY6G (C) immunofluorescence in small intestines from control and TNBS-treated mice. (D) MPTX2 immunofluorescence in small intestines from control and TNBS-treated mice. (E) Representative H&E-stained small intestines from TNBS-treated $Lsr^{ctrl}$ and $Lsr^{vill\ KO}$ mice. (F and G) F4/80 (F) and LY6G (G) immunofluorescence in small intestines from TNBS-treated $Lsr^{ctrl}$ and $Lsr^{vill\ KO}$ mice. (H) Representative H&E-stained small intestines from TNBS-treated mice administered with AAV-CTL or AAV-Lsr. (I and J) F4/80 (I) and LY6G (J) immunofluorescence in small intestines from TNBS-treated mice administered with AAV-CTL or AAV-Lsr. Scale bars: A, E and H, 100 μm; B, C, D, F, G, I, and J, 50 μm.

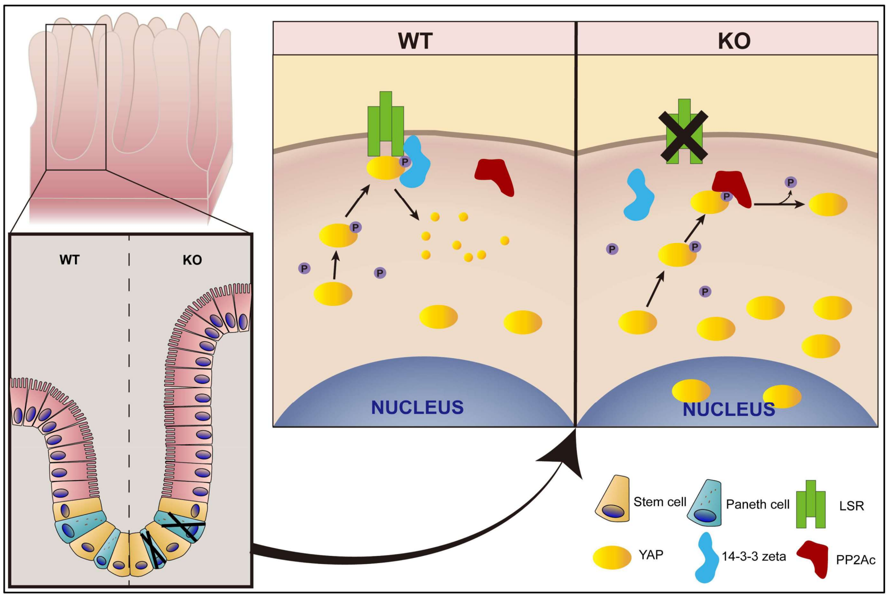


**Supplementary Figure 5. Summary.**

In wildtype intestine (WT), LSR interacts with 14-3-3 zeta and YAP, thus facilitating 14-3-3-dependent YAP ubiquitination and degradation by the proteasome pathway. ISCs differentiate into Paneth cells and other epithelial cells to form the intestinal barrier and maintain the intestinal homeostasis. When LSR was deleted (KO), PP2Ac competitively bound to YAP and dephosphorylated it, thus inhibiting its degradation and translocating it into the nucleus. Enhanced abundance and nuclear localization of YAP in mouse intestinal epithelium resulted in increased numbers of proliferating cells and loss of Paneth cell lineage.
