## Supplementary Table1 for "LSR Targets YAP to Modulate Intestinal Paneth Cell Differentiation"

**Supplementary Table 1: primary antibody list**

| Target | Product name | Host | Applications | Cat. NO. | Company |
|---|---|---|---|---|---|
| LSR | LSR Purified MaxPab mouse Polyclonal Antibody (B01P) | Mouse | IF, WB | H00051599-B01P | Abnova |
| LSR | LSR (D3E3N) XP Rabbit mAb | Rabbit | CoIP | #14804 | Cell Signaling Technology |
| LSR | LSR / LISCH7 Rabbit anti-Mouse Polyclonal (aa35-205) Antibody | Rabbit | IF | LS-C373180 | LifeSpan Biosciences |
| YAP | YAP1 Antibody (63.7) | Mouse | IF, WB | sc-101199 | Santa Cruz Biotechnology |
| YAP | YAP (D8H1X) XP Rabbit mAb | Rabbit | CoIP | #14074 | Cell Signaling Technology |
| p-YAP | Phospho-YAP(Ser397) (D1E7Y) Rabbit mAb | Rabbit | WB | #13619 | Cell Signaling Technology |
| MPTX2 | Rabbit monoclonal [EPR20920-19] to MPTX+MPTX2 | Rabbit | IF | Ab238123 | Abcam |
| LYZ | Lysozyme Recombinant Rabbit Monoclonal Antibody (ST50-02) | Mouse | IF | MA5-32154 | Invitrogen |
| OLFM4 | Olfm4 (D6Y5A) Rabbit mAb (Mouse Specific) | Rabbit | IF | #39141 | Cell Signaling Technology |
| OLFM4 | Olfm4 (D1E4M) Rabbit mAb | Rabbit | IF | # 14369 | |
| LGR5 | LGR5 Monoclonal Antibody (OTI2A2) | Mouse | IF | MA5-25644 | Invitrogen |
| LGR5 | LGR5 Mouse Monoclonal Antibody [Clone ID: UMAB212] | Mouse | WB | UM800104 | Origene |
| EpCAM | EpCAM (E8Q1Z) Rabbit mAb (Mouse Preferred) | Rabbit | IF | #42515 | Cell Signaling Technology |
| Ubiquitin | Ubiquitin Recombinant Rabbit | Rabbit | WB | 701339 | Invitrogen |

| | | | | | |
|---|---|---|---|---|---|
| | Monoclonal Antibody (10H4L21) | | | | |
| PPP1ca | PPP1A (PPP1CA) Rabbit Polyclonal Antibody | Rabbit | CoIP, WB | TA327322 | Origene |
| PP2ca | PP2AC Subunit Antibody | Rabbit | CoIP, WB | #2038 | Cell Signaling Technology |
| 14-3-3 ζ/δ | 14-3-3 ζ/δ (D7H5) Rabbit mAb | Rabbit | CoIP, WB | #7413 | Cell Signaling Technology |
| 14-3-3 γ | 14-3-3 γ (D-6) | Mouse | CoIP, WB | sc-398423 | Santa Cruz Biotechnology |
| PCNA | PCNA Monoclonal Antibody (PC10) | Mouse | IF, WB | MA5-11358 | Invitrogen |
| Ki67 | Anti-Ki67 antibody [SP6] | Rabbit | IF, WB | ab16667 | Abcam |
| F4/80 | F4/80 Monoclonal Antibody (BM8), eBioscience | Rat | IF | 14-4801-85 | Invitrogen |
| LY6G | Purified anti-mouse Ly-6G Antibody (1A8) | Mouse | IF | 127602 | Biolegend |
| ZO-1 | ZO-1 Polyclonal Antibody | Rabbit | IF | 61-7300 | Invitrogen |
| Tricellulin | MARVELD2 Polyclonal Antibody | Rabbit | IF | 48-8400 | Invitrogen |
| CLDN1 | Claudin 1 Monoclonal Antibody (2H10D10) | Mouse | IF | 37-4900 | Invitrogen |
| CLDN4 | Claudin 4 Polyclonal Antibody (ZMD.306) | Rabbit | IF | 36-4800 | Invitrogen |
| Anti-Active-β-Catenin | Anti-Active-β-Catenin (Anti-ABC) Antibody, clone 8E7 | Mouse | IF | 05-665 | Sigma-Aldrich |
| SOX9 | Anti-SOX9 antibody [EPR14335] | Rabbit | WB | ab185230 | abcam |
| ATOH1 | ATOH1 Polyclonal antibody | Rabbit | WB | 21215-1-AP | Proteintech |
| P-Lats1 | Phospho-LATS1 (Thr1079) Antibody | Rabbit | WB | AF7169 | Affinity |
| β-actin | Anti-beta Actin Antibody - | Mouse | WB | ab8226 | Abcam |

| | |
|---|---|
| | [mAbcam 8226]<br>Loading Control |
