## Supplementary Table2 for "LSR Targets YAP to Modulate Intestinal Paneth Cell Differentiation"

**Supplementary Table 2: Primer sequences used for qRT-PCR**

| Gene | Forward | Reverse |
|---|---|---|
| Lsr | CTACAACCCCTATGTGGAGTGC | CTGCCCTGGTAGTAGTCTCCC |
| Lgr5 | AGAGCCTGATACCATCTGCAAAC | TGAAGGTCGTCCACACTGTTGC |
| Lyz1 | TGACTCTGGGACTCCTCCTG | CTTGACACCACGGTAGCCAT |
| Itln1 | TCCAGTCAGCAAGGCAACAGAG | CAGGTTCTCAGCCTGGATGTCA |
| Mptx2 | AGTGTAGCACAAACAGACATGAA | TCAAGCCTTTTCTCCCAAATGC |
| Defa5 | GAAGAGGACCAGGCTGTGTC | GGTCCCAAAAACGCGTTCTC |
| Cd24a | ACATCTGTTGCACCGTTTCCCG | CAGGAGACCAGCTGTGGACTG |
| Cdx1 | CACAGAGCGGCAGGTAAAGA | TAGACTTCCTAGGGGTGGCC |
| Cdx2 | GCAGTCCCTAGGAAGCCAAG | GCCAGCTCACTTTTCCTCCT |
| Muc2 | CCTGAAGACTGTCGTGCTGT | GGGTAGGGTCACCTCCATCT |
| Spdef | AAGGCAGCATCAGGAGCAAT | TCAATGACGGGACACTGCTC |
| Nuerog3 | CGAAGCAGAAGTGGGTGACT | CGATCATTGGCCTTCTTGCG |
| Chga | GAAGTGCGTCCTGGAAGTCA | CCTTGGAGAGCCAGGTCTTG |
| Yap | TGAGATCCCTGATGATGTACCAC | TGTTGTTGTCTGATCGTTGTGAT |
| Ctgf | AGAACTGTGTACGGAGCGTG | CTTTGGAAGGACTCACCGCT |
| Edn1 | GGAGCTCCAGAAACAGCTGT | GCCTGGTCTGTGGCCTTATT |
| Cry61 | CGCAATTGGAAAAGGCAGCT | CGATGCATTTCTGGCCATGG |
| Il1α | ACGTCAAGCAACGGGAAGAT | ACTTCTGCCTGACGAGCTTC |
| Il6 | GAGGATACCACTCCCAACAGACC | AAGTGCATCATCGTTGTTCATACA |
| Il2 | GAGCAGCTGTTGATGGACCT | AATCCAGAACATGCCGCAGA |
| Tnfα | CGTCGTAGCAAACCACCAAG | TTGAAGAGAACCTGGGAGTAGACA |
| Ifnγ | GCAAGGCGAAAAAGGATGCA | CGACTCCTTTTCCGCTTCCT |
| CD44 | ATCCTCGTCACGTCCAACAC | GCTTTCTGGGGTGCTCTTCT |
| Ccnd1 | CCCTGGAGCCCTTGAAGAAG | AGATGCACAACTTCTCGGCA |
| Axin2 | CCTGACCAAACAGACGACGA | CACCTCTGCTGCCACAAAAC |
| Defa2 | CACTTGTCCTCCTCTCTGCC | GAGACAGACACAGCCTGGTC |
| Defa3 | GACCAGGCTGTGTCTGTCTC | CCCTTTCTGCAGGTCCCATT |
| Defa17 | GACCAGGCTGTGTCTGTCTC | CCCTTTCTGCAGGTCCCATT |

| | | |
|---|---|---|
| Defa20 | CACTTGTCCTCCTCTCTGCC | GAGACAGACACAGCCTGGTC |
| Defa21 | CTCCTCTCTGCCCTCATCCT | GAGACAGACACAGCCTGGTC |
| Defa22 | GAAGAGGACCAGGCTGTGTC | CTCCTCTATTGCAGCGACGT |
| Defa23 | GACCAGGCTGTGTCTGTCTC | CCCTTTCTGCAGGTCCCATT |
| Defa24 | GAAGAGGACCAGGCTGTGTC | GCGTTCTCTTCCTTTGCAGC |
| Defa26 | GAAGAGGACCAGGCTGTGTC | TGCAGGTCCCATTAATGCGT |
| Defa28 | TGATGAGCAGCCAGGGAAAG | ATCCGATCTCTCAACGCGTC |
| Defa29 | CTTCCAGGTCCAGGCTGATC | CTTCGCACAATGGCTGCTTT |
| Defa30 | GAAGAGGACCAGGCTGTGTC | GCGTTCTCTTCCTTTGCAGC |
| Defa31 | GACCAGGCTGTGTCTGTCTC | CCCTTTCTGCAGGTCCCATT |
| Defa32 | CACTTGTCCTCCTCTCTGCC | GAGACAGACACAGCCTGGTC |
| Defa33 | CACTTGTCCTCCTCTCTGCC | GAGACAGACACAGCCTGGTC |
| Defa35 | GAAGAGGACCAGGCTGTGTC | GGTCCCAAAAACGCGTTCTC |
| Defa37 | GAAGAGGACCAGGCTGTGTC | GGTCCCAAAAACGCGTTCTC |
| β-actin | CATTGCTGACAGGATGCAGAAGG | TGCTGGAAGGTGGACAGTGAGG |
| LSR-sRNA | TTTGAAGGAACACTGATGA | |
| AAV-m-Lsr | Ttgacctccatagaagacaccgggatcc | Tgtagtcgttaattaaggtac |
| | AAGCTTGCCACCATGGCGCC | CGAATTCGACGACTAAACTTTCCCGACT |
